# Extracellular matrix defects destabilise apical cytoarchitecture and mechanical properties during early Down syndrome neurodevelopment

**DOI:** 10.64898/2026.06.22.733824

**Authors:** Teresa P. Silva, Ekaterina R. Iijima, Ioakeim Ampartzidis, Grace Cooksley, Blanca Sanchez Paya, Ekin Ucuncu, Daniel Holder, Imogen Smith, James Smith, Giovanni Giuseppe Giobbe, Paolo De Coppi, Kathleen J. Millen, Jonathan D. W. Clarke, Frederick J. Livesey, Nicholas D. E. Greene, Parthiv Haldipur, Gabriel Galea, Paula Alexandre

## Abstract

Down syndrome (DS) is associated with altered brain development, especially in the cerebellum. However, how trisomy 21 (TS21) is linked to cerebellar defects remains poorly understood. Cerebellar organoids reveal that TS21 leads to extracellular matrix (ECM) alterations, perturbing downstream pathways linked to reduction of apical RAB11^+^ endosomes. These changes lead to impaired apical maintenance and altered progenitor composition. Significantly, ECM-enriched culture rescues apical defects in TS21 organoids, supporting a functional role for ECM in preserving apical organisation and progenitor niche integrity. In the developing human DS cerebellum and 2D neural cultures, where exogenous ECM is present, severe epithelial disorganisation is attenuated. Nevertheless, ECM organisation and mechanical properties remain altered in 2D cultures. Our findings identify TS21-driven ECM abnormalities as a mechanism impairing apical integrity, progenitor composition and mechanical proprieties in neural stem cell models of DS. Altogether, we established apical maintenance instability as an early developmental defect contributing for DS neuropathology.

## Introduction

Down syndrome (DS) is one of the most common chromosomal abnormalities caused in 95% of cases by three free copies of chromosome 21 (HSA21), referred as trisomy 21 (TS21) ^1^. Individuals with DS present with an average brain volume reduction ^2,3^. Although most of previous DS studies have primarily focused on cortical phenotype, it is the cerebellum that shows the largest reduction in volume ^2^. Human DS foetuses (17-21 weeks of gestation) present with a significant decrease of rhombic lip and granule cell proliferation in the cerebellum^4^, which suggests early developmental defect. This is accompanied by a significant volume reduction and simplified folding pattern ^2–4^. However, the underlying mechanisms leading to decreased cell proliferation and consequent reduction in cerebellar size remain poorly understood.

In the neural tissue, extracellular matrix (ECM) components localise at the basal surface of the neuroepithelium. These provide structural support and regulate cell signalling involved in cytoskeleton remodelling, and maintenance of neuroepithelial cell polarity, cell survival and proliferation, and tissue mechanics ^5,6^. During cortical development, apical and basal radial glia progenitor types extend basal processes that contact the ECM on the pial surface. ECM receptors are essential for anchoring radial glia basal processes to the basement membrane, and these signalling interactions are critical to promote their survival and self-renewal ^7^. In mice lacking integrin β1 or laminin α2/4, radial glial cells lose contact with pial surface, leading to increased apoptosis and microcephaly ^7^. Similarly, inhibition of integrin signalling in ferrets reduces basal radial glial populations ^8^, potentially due to impaired self-renewal ability and survival.

ECM defects have been previously reported using different models of DS. ECM-related genes are found upregulated in mouse DS embryonic heart tissue ^9^ and fibroblasts ^10^, including genes mapping to HSA21 (*ADAMTS1, ADAMTS5, COL6A1, COL6A2* and *COL18A1*). This up-regulation was shown to drive alterations in tissue mechanics and mechano-transduction during heart development ^9^. In the human brain, DS cortical tissue exhibited a delayed folding response to the ECM components HAPLN1, lumican, and collagen I (HLC) relative to controls ^11^. Moreover, single cell multiomic analysis of human developing neocortex showed a significant alteration in vascular ECM gene expression and radial glia integrin signalling, together with a reduction in neural progenitors and accelerated neuronal differentiation in DS^12^. This suggests DS-derived ECM dysregulation could explain the simplified gyrification pattern, growth and proliferation defects ^2,4,6^ observed in DS brains. However, the link between ECM and DS brain phenotype remains largely unexplored.

Human TS21-derived induced pluripotent stem cells (iPSCs) provide opportunities to investigate the mechanisms underlying altered brain development in DS ^13^. TS21 iPSC-derived astrocytes show a global alteration in chromatin state and gene expression of cell adhesion and ECM genes ^14^. In addition, a recent analysis of OLIG2, a regulator of oligodendrocyte differentiation encoded by HSA21 ^15^, shows an increased proportion of OLIG2^+^ progenitor cells in TS21-derived iPSC differentiated into 2D cultures and organoids ^16,17^. When TS21 iPSC are differentiated into neurons, synapses are formed albeit with impaired function in both culture following xenotransplantation into mouse brain ^18^. Reduced excitatory synaptic density and functional deficits in excitatory synaptic transmission were also found in human TS21 iPSC-derived neurons. This suggests disrupted synapse development in DS ^19,20^. In TS21 cortical organoids, a neurogenesis defect has been reported, characterised by reduced progenitor proliferation and neuronal production ^21^. On the other hand, choroid plexus organoids differentiated from TS21 iPSC show no difference in organoid growth rates ^22^. However, they do exhibit defects in ciliogenesis and epithelial cell polarity, together with global gene expression alteration of nervous system development, cilium movement and motility, cell adhesion, and basement membrane genes ^22^. It is unknown if these changes occur in the cerebellum and whether TS21-related ECM defects affect apical organisation and progenitor proliferation.

In this study, we have employed cerebellar organoids ^23^and neuroepithelial sheets ^24,25^ derived from human disomy and TS21-derived iPSCs. We have also performed histological analysis of human foetuses, to investigate the impact of TS21-derived defects on the DS brain development. We observed that ECM defects in TS21 can impair apical junction maintenance and alter progenitor cell type composition. These phenotypes can then be rescued by supplementation with an exogenous source of ECM - Matrigel. ECM downstream signalling is also affected and reduction of RAB11^+^ endosomes at the apical surface likely contribute to the apical junction loss. Although less severe apical defects are present in human tissue and 2D cultures, ECM localisation and mechanics are still significantly altered in iPSC-derived 2D cultures. Our investigation uncovers TS21 and ECM-derived alterations in apical maintenance, proliferation and mechanical properties in TS21 iPSC-derived neural models. Altogether, we demonstrate that defective apical maintenance is an early neurodevelopmental feature of Down syndrome.

## Results

### 1. Altered proportion of apical-basal mitotic progenitors in TS21 cerebellar organoids

To model TS21-related defects in DS brain tissue, human iPSC-derived cerebellar organoids were generated (**Figure 1A**) ^23^, as cerebellum is one of the most severely affected brain structures in DS ^2^. For that, three control iPSC lines, two full TS21 iPSC lines, and one partial TS21 iPSC line were used (**Figure S1 and supplementary table S1**). The resulting self-organising cerebellar organoids develop neuroepithelial structures that contain an apical junctional belt expressing apical polarity and junctional proteins, including F-actin, tight-junction protein ZO1, adherens junction N-cadherin (NCAD), that frequently outline a lumen (**Figure S2A**). These structures contain neuroepithelial cells expressing proliferative markers SOX2, phospho-vimentin (pVIM), and KI67 (**Figure S2B**). In addition, TUJ1^+^ and HuC/D^+^ neurons can be found surrounding the neuroepithelium (**Figure S2B**). Sparse labelling of neuroepithelial cells with viral GFP demonstrated their elongated morphologies spanning from apical surface to the basal lamina (**Figure S2C**). The basal end-feet of these neuroepithelial cells can contact the surrounding laminin, located on the basal region of the neuroepithelium. Both disomy and TS21-derived organoids exhibited ATOH1^+^ and SKOR2^+^ cerebellar progenitors (**Figure S2D**).

**Figure 1.**
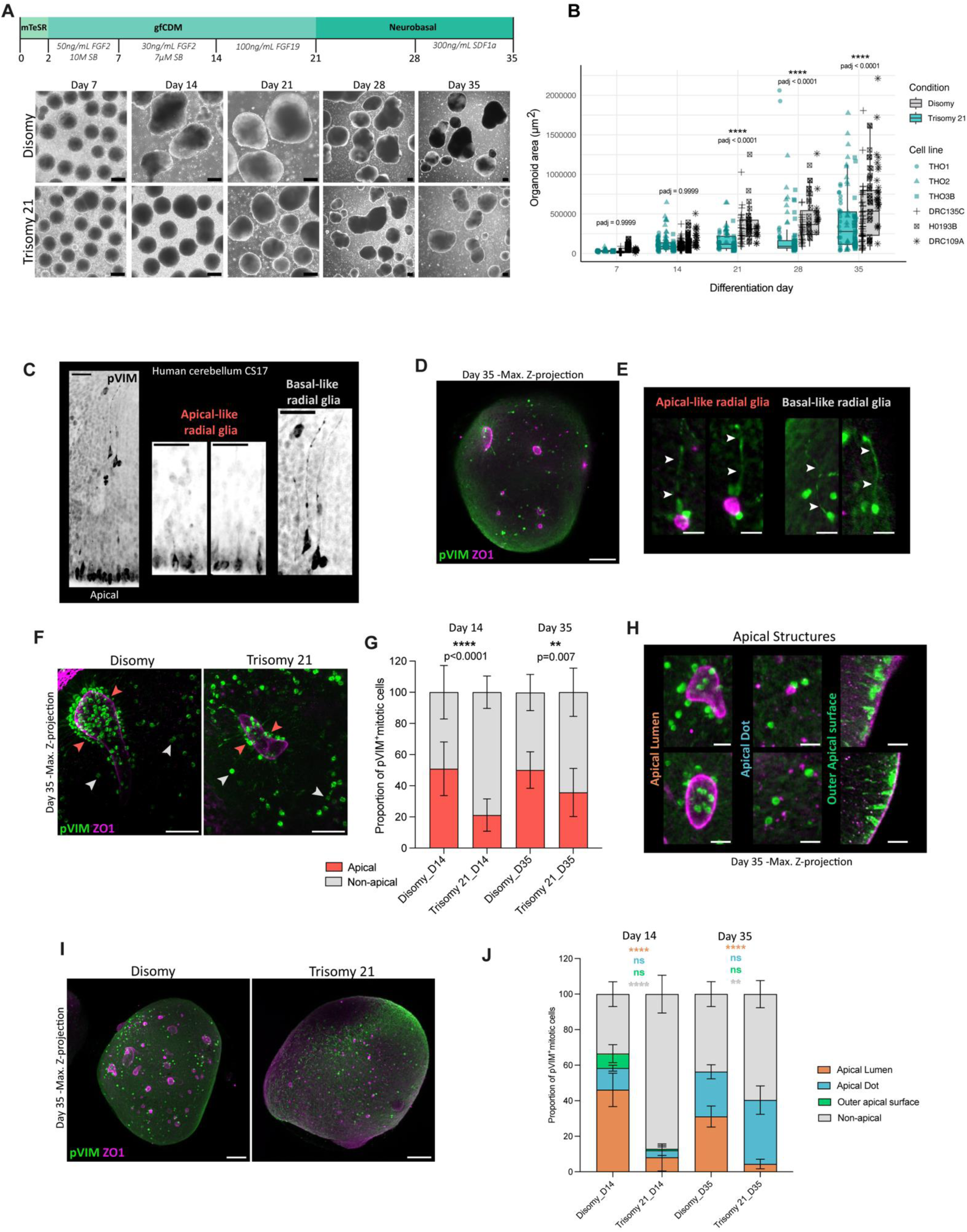
Imbalance of apical-basal proliferation in trisomy 21 (TS21) organoids. **A.** Schematic representation of human iPSC differentiation into cerebellar organoids with representative brightfield images across cerebellar differentiation, showing organoid cultures from disomy and TS21 conditions. Scale bar, 200μm. **B.** Area of human iPSC-derived organoids across differentiation. Box-plot shows the distribution of organoid area for each condition at each time point, with individual organoids shown as points. Symbol shapes indicate independent cell lines. Data from 3 TS21 and 3 disomic iPSC lines. Two-way ANOVA with Tukey’s multiple comparisons test. **C.** Human cerebellum immunostaining for pVIM at CS17, with zoom-in showing both apical and basal glia-like cells. Scale bar, 20μm. **D.** Representative maximum Z-projection of whole mount immunofluorescence images showing apical ZO1 expression and pVIM-expressing mitotic progenitors in control disomic organoids at day 35. Scale bar, 100 μm. **E.** High magnification of pVIM^+^ cells near to apical (ZO1+) and non-apical (ZO1-) pVIM^+^ regions, resembling apical and basal radial glia-like cells, respectively. The arrowheads indicate the projections in radial glia-like cells. Scale bar, 20 μm. **F.** Whole mount immunofluorescence images for pVIM and ZO1. The arrowheads indicate pVIM expressing progenitors nearby apical lumens and non-apical regions in red and grey, respectively. Scale bar, 50 μm. **G.** Stacked bar diagrams depict the proportion of pVIM^+^ cells proliferating near to apical or non-apical regions at days 14 and 35. Data obtained from 3 disomic and 3 TS21 cell lines. Two-way ANOVA (Uncorrected Fisher’s LSD); error bars represent SEM. **p = 0.007, ****p < 0.0001. **H.** Representative immunofluorescence images of pVIM cells close to distinct apical ZO1 structures, including lumen, dot and outer apical surfaces. Scale bar, 20 μm. **I.** Representative maximum Z-projection of whole mount immunofluorescence images of ZO1 and pVIM at day 35. Scale bar, 100 μm. **J.** Stacked bar graph shows the proportion of pVIM^+^ mitotic cells associated with apical structures – lumen, dot and outer apical surface – and in non-apical regions. Data obtained from 1 control (H0193B) and 1 TS21 (THO3B) cell line. Error bars represent SEM. Two-way ANOVA followed by Šídák’s multiple comparisons test. **Disomy D14 vs D35:** Apical lumen-like **, Apical dot-like *, Outer apical surface – ns, Non-apical - ns; **Trisomy D14 vs D35:** Apical lumen-like - ns, Apical dot-like ****, Outer apical surface - ns; Non-apical ****. ns, not significant; *p < 0.05, ** p < 0.01, ****p < 0.0001.

To analyse the cerebellar organoids growth, we quantified organoid size and cell proliferation at different timepoints (**Figures 1A and 1B**). Cerebellar organoid area increased significantly over time in both conditions (**Figure S2E**), but control organoids became significantly larger than TS21 organoids from day 21 onwards (**Figure 1B**). However, no differences in overall proliferation were found in TS21-derived organoids when compared with disomic controls (**Figures S2F, S2G and S2H**), which suggests similar growth rates. Cerebellar primary progenitor zones containing apical and basal progenitor populations have been already associated with expansion of human developing cerebellum ^26^. Human cerebellum contains mitotic pVIM^+^ progenitors in both ventricular (VZ) and subventricular (SVZ) zones with long radial processes resembling apical and basal radial glia ^26^ (**Figure 1C**). At day 35, 3D whole-mount staining of control iPSC-derived organoids for ZO1 and pVIM markers (**Figure 1D**) revealed mitotic cells with a single long process in both apical (ZO1^+^) and non-apical (ZO1^-^) positions (**Figure 1E**), which resemble the presence of two types of radial glia-like cells as seen in the human cerebellum. 3D whole-mount staining for ZO1 and pVIM was performed in cerebellar organoids at days 14 and 35 to determine the proportion of progenitors dividing at and away from apical surfaces (**Figure 1F**). The proportion of apical versus non-apical mitoses was significantly different in TS21-derived cerebellar organoids, with a significant reduction in the proportion of apical mitotic progenitors, when compared to disomic controls at days 14 and 35 (**Figure 1G**). For this apical quantification, all mitotic cells in contact with a ZO1^+^ apical structure were considered, including lumen, dot and outer apical surface (**Figure 1H**). Further to this, we noticed an enrichment of dot-like ZO1^+^ inner apical surfaces in the TS21 organoids relative to the disomy (**Figure 1I**). Thus, we quantified mitotic cells by their contact with a ZO1^+^ apical structure subdivided into lumen, dot and outer apical surface, as well as mitotic non-apical progenitor populations in TS21 (THO3B) and control (HO193B)-derived organoids. At day 35, most of the apical mitoses in TS21-derived organoids were located near to an apical dot, while a small proportion was found attaching an apical lumen. A significant increase of mitoses near apical dot was observed when compared to day 14, consistent with the enrichment of dot-like apical structures in these organoids (**Figures 1I and 1J**). This suggests that alterations on apical structures may contribute to an imbalance between apical and non-apical progenitor populations.

### 2. Apical structures are reduced in TS21-derived cerebellar organoids

To investigate whether alteration on progenitor cell type composition is associated with a defective apical surface, immunostaining for ZO1 was performed to outline the apical surface in cerebellar organoids at day 14. In the TS21-derived organoid cryosections, we observed that almost half of the organoids did not have open apical lumens by day 14 (**Figure S3A and S3B**). Thus, to maximise the chances of identifying lumens in TS21, we performed whole mount organoid immunostaining analysis. Apical lumens expressing ZO1 were identified in both control and TS21 cerebellar organoids by day 14 (**Figure 2A**). Then, the volume of the five largest apical lumens per organoid was manually determined (**Figure 2B**), revealing that TS21-derived organoids exhibited a significantly smaller lumen volume (**Figure 2C**). To determine whether the reduction of lumen volume in TS21 could be associated with alterations in organoid volume, correlation analysis between lumen and total organoid volumes at day 14 was performed (**Figure 2D**). A positive correlation was observed only in the controls (**Figure 2D**). Furthermore, when lumen volume was normalised to the total organoid volume, a significant reduction was also observed in TS21 organoids (**Figure 2E**). In addition to apical lumens, ZO1 expression was also detected as apical puncta and at the outer surface of the organoids (**Figure 2F**). To quantify the proportion of organoid volume expressing ZO1, we separated ZO1^+^ regions into inner apical structures, including apical puncta and lumens, and outer ZO1^+^ surfaces (**Figure 2F**). For this, a Fiji-based segmentation pipeline was used to “peel” the organoids into inner and outer apical surfaces, and the volume of ZO1^+^ signal was then normalised to the total organoid volume. At day 14, TS21 organoids showed an overall significant reduction in ZO1 expression at both inner and outer surfaces (**Figure 2G**).

**Figure 2.**
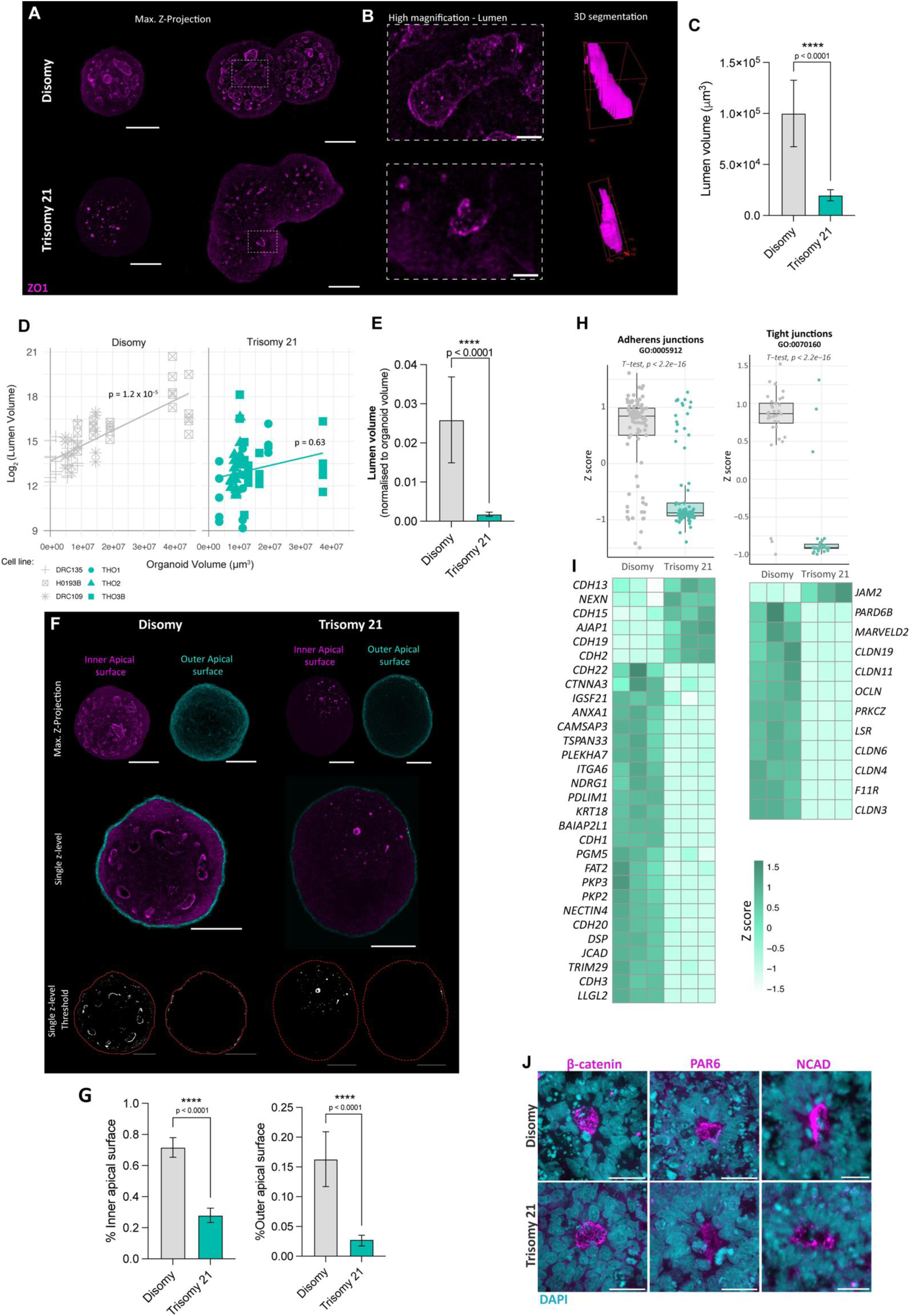
TS21 organoids show apical defects at day 14. **A.** Whole mount immunofluorescence staining of ZO1 showing apical lumens in disomic and TS21 conditions at day 14. Scale bar, 100 μm. **B.** Representative high magnification of apical lumens and 3D segmentation for lumen size quantification. Scale bar, 20 μm. **C.** Bar-plot compares the volume of the five largest ZO1^+^ apical lumens per organoid in disomy and TS21 at day 14. Data obtained from 3 disomic and 3 TS21 cell lines. Mann-Whitney test; error bars represent SEM. ****p < 0.0001. **D.** Scatter plot of organoid volume against log₂ (lumen volume) in disomic and TS21 organoids at day 14. Each point represents an individual lumen, and different symbols indicate independent cell lines. Linear regression lines are shown for each condition. Lumen volume scaling significantly with total organoid volume in disomy (Pearson r = 0.54; p = 1.2 x 10^-5^), but no significant relationship in TS21 condition (Pearson r = 0.07; p = 0.63). Data include the five largest ZO1^+^ apical lumens, obtained from 3 control (H0193B) and 3 TS21 (THO3B) cell lines. **E.** Bar-plot depicts the five largest lumen volumes normalised to total organoid volume. Data obtained from 3 disomic and 3 TS21 cell lines. Mann-Whitney test; error bars represent SEM. ****p < 0.0001. **F.** Representative whole mount immunofluorescence images showing ZO1 segmentation to split inner and outer apical surfaces, with respective thresholds used to quantify the percentage of inner and outer apical surface. Red dash outlines the organoid. Scale bar, 100 μm. **G.** Bar-plots comparing the percentage of inner and outer apical surface positive for ZO1 in disomy and TS21 organoids. Data obtained from 3 disomic and 3 TS21 cell lines. Mann-Whitney test; error bars represent SEM. ****p < 0.0001. **H.** Box blot showing distribution of genes annotated in adherens (GO:005912) and tight (GO:0070160) junctions. Data were obtained from bulk RNAseq and represent minimum to maximum Z-score. **I.** Heatmap highlighting genes related to adherens and tight junctions from bulk RNAseq data. Values are shown as Z-score. Data obtained from 3 differentiation for control (H0193B) and 3 TS21 (THO3B) cell lines. **J.** Immunofluorescence staining for apical markers, including β-catenin, PAR3 and NCAD in organoid cryosections at day 14. Scale bar, 20 μm.

To determine whether the disruption of apical components also occurred at the transcriptional level, bulk RNA-sequencing (RNAseq) analysis of cerebellar organoids derived from one TS21 (THO3B) and one control (HO193B) cell line was performed by day 14. A significant number of genes were found differentially expressed in TS21 condition (**Figure S3C**). When analysed by chromosome expression, the ratio of differential expressed genes per total number of genes per chromosome showed a higher proportion in HSA21 (**Figure S3D**), consistent with HSA21 triplication genotype. Consistent with ZO1 staining analyses, bulk transcriptomic analysis also revealed that the significantly down-regulated genes in TS21 organoids were highly associated with apical-related cellular components, including tight junction (GO:0070160) and adherens junction (GO:005912; **Figure S3E**). Further analysis of the Z-score for differentially expressed genes annotated in both tight and adherens junctions GO terms confirmed a significant decrease of these apical transcripts in TS21 organoids when compared with controls (**Figure 2H**). Among these transcripts, we identify several genes important for cell adhesion, including claudins (*CLDN19, 11, 6,* and *4*), *OCLN*, *CDH3, CDH1, F11R* and *ITGA6*, but also to in anchoring junction subnetworks, including *PLEKHA7* and *CAMSAP3* (involved in anchoring microtubules to zonula adherens) and *PDLIM1* (an adapter that brings other proteins to the cytoskeleton), suggesting a disrupted apical junctional network (**Figure 2I**). Surprisingly, the neuroepithelial structures formed in TS21 cerebellar organoids still accumulated apical proteins at the luminal apical surface, including β-catenin, PAR6 and NCAD (**Figure 2J**). These findings suggest that TS21-derived neuroepithelial cells retain the ability to polarise, but the induction or maintenance of apical luminal structures may be impaired in TS21 cerebellar organoids.

### 3. Ability for apical expansion and maintenance is disrupted in TS21-derived cerebellar organoids

To elucidate the mechanisms underlying lumen formation in our disomy cerebellar organoids and potential defects in TS21 organoids, we differentiated a disomic ZO-1 mEGFP (ZO1-GFP) iPSC line into cerebellar organoids and performed daily time-lapse imaging from day 2 to day 14, acquiring images every 20 minutes over periods of more than 14 hours. At day 2, ZO1-GFP iPSC-derived aggregates showed signs of early polarisation (**Figure 3A**). A few hours after neural induction, apical reorganisation was observed (**Figure S4A**), followed by changes in apical junction morphology and size as shown at day 4 (**Figure S4B**). From day 3, ZO1- GFP organoids formed a continuous ZO1-GFP at the outer apical surface (**Figure 3A**). As the organoid development progressed, the apical surface area of individual cells decreased until day 6 (**Figure 3B**), resulting in a more homogeneous honeycomb-like pattern (**Figure 3A**, **3B and S4C**). Apical luminal surfaces (identified by the expression of ZO1-GFP inside the organoid i.e. inner ZO1-GFP) were also observed in the cerebellar organoids from day 2 to day 6 (**Figure S4D**).

**Figure 3.**
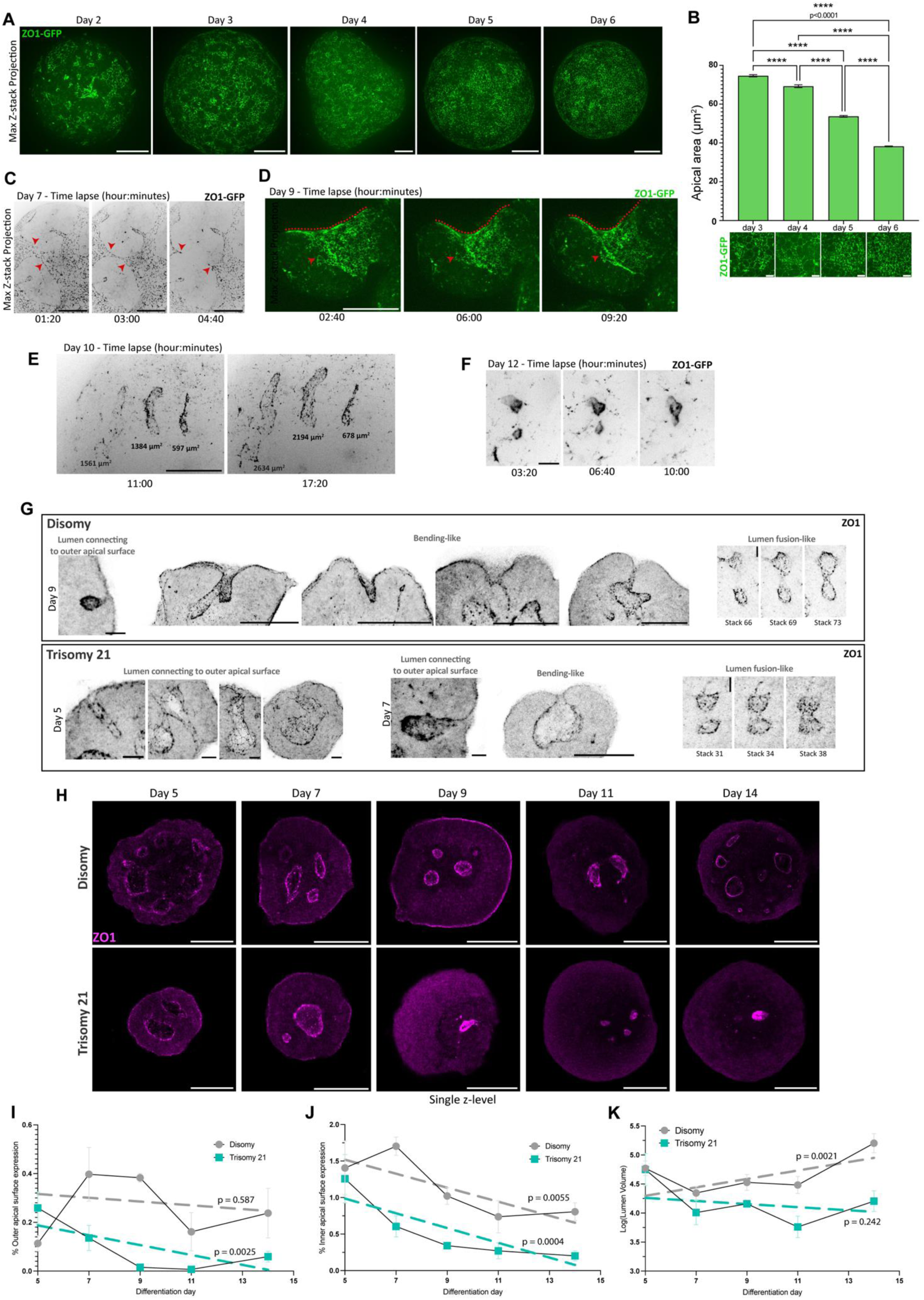
Lumen formation and apical dynamics in cerebellar organoids. **A.** Representative maximum intensity Z-projection of disomic ZO1-GFP organoids from day 2 to 6 of differentiation. Scale bar, 100 μm. **B.** Bar-plot showing the mean apical surface area per cell in ZO1-GFP organoids between day 3 and 6, with representative maximum intensity Z-projection images of apical surfaces for each timepoint. Scale bar, 20 μm. Data obtained from ≥ 5 organoids per timepoint from 2 differentiations. One-way ANOVA (Kruskal-Wallis test); error bars represent SEM. ****p < 0.0001. **C.** Image sequence time-lapses from ZO1-GFP organoids at day 7, with arrowheads indicating apical surface breaks. Scale bar, 100 μm. **D.** Representative time-lapse images of day 9 ZO1-GFP organoids suggesting bending of the organoids, indicated by dashed red lines and arrowheads. Scale bar, 100 μm. **E.** Time-lapse frames of live imaging of ZO1-GFP-derived cerebellar organoids at day 10, showing lumen growth. Scale bar, 100μm. **F.** Image sequence from time-lapses of day 12 ZO1-GFP organoids, demonstrating fusion of lumens. Scale bar, 20μm. **G.** Representative maximum Z-projection from whole mount immunofluorescence staining for ZO1, suggesting lumen connection with outer apical surface (scale bar, 20 μm), bending-like (scale bar, 100 μm), and lumen fusion (scale bar, 20 μm) in disomy and TS21 conditions. **H.** Representative single Z-level from whole mount immunofluorescence staining for ZO1, showing apical lumens in disomy and TS21 conditions between day 5 and 14. Scale bar, 100 μm. **I.** Line plot showing % outer apical surface expressing ZO1 over differentiation timepoint. Points represent mean value at each time point, with error bars representing SEM. Dashed lines represent linear regression analysis for each condition. Disomic controls show no significant relationship between outer apical surface and timepoint (slope = -0.0075, 95% CI: -0.037 to 0.022, R² = 0.018, p = 0.587). In contrast, TS21 organoids show a significant decrease in apical surface with increasing time (slope = -0.0203, 95% CI: -0.032 to -0.008, R² = 0.407, p = 0.0025). Data obtained from one disomy (H0193B) and one TS21 (THO3B) cell line. **J.** Line plot shows % inner apical surface across timepoints. Points show mean at each time point, with error bars representing SEM. Dashed lines represent linear regression analysis for each condition. Both disomic and TS21 organoids showing significant negative associations, indicating a progressive decrease over time in both conditions: disomy, slope = −0.0959, 95% CI: -0.160 to -0.032, R² = 0.373, p = 0.0055; TS21, slope = -0.1010, 95% CI: -0.150 to -0.052, R² = 0.513, P = 0.0004. Data obtained from one disomy (H0193B) and one TS21 (THO3B) cell line. **K.** Line plot showing log₂ (lumen volume) across differentiation days in disomic and TS21 organoids. Points represent mean lumen volume at each time point, with error bars indicating SEM. Dashed lines represent linear regression analysis for each condition. Disomic condition shows a significant positive association (slope = 3.89 × 10⁴, 95% CI: 1.46 × 10⁴ to 6.32 × 10⁴, R² = 0.124, p = 0.0021), whereas TS21 shows no significant association (slope = -3.62 × 10³, 95% CI: -9.77 × 10³ to 2.54 × 10³, R² = 0.035, p = 0.242), differing significantly between conditions. (Z = 3.39, p < 0.001). Data include the five largest ZO1^+^ apical lumens per organoid, obtained from 1 control (H0193B) and 1 TS21 (THO3B) cell line.

By day 7, discontinuities the outer ZO1-GFP regions became detectable and persisted until day 13 (**Figures 3C and S4E**), coinciding with pronounced changes in organoid shape. Bending of the outer surface was present by day 9 (**Figure 3D**), a process that resembles the tissue movements of mammalian neural tube closure ^27^ and reported in iPSC-derived spinal cord organoids ^28^. The outer apical surface progressively internalised at day-13 timelapses (**Figure S4F**). In addition, we also observed ZO1-GFP-labelled apical luminal expansion and fusion at days 10 and 12 (**Figures 3E and 3F**), suggesting that several apical remodelling mechanisms can act simultaneously to generate luminal apical surfaces in the cerebellar organoids.

To investigate whether the disruption of apical structures in TS21 cerebellar organoids resulted from morphogenetic defects, we analysed ZO1-stained organoids differentiated from one TS21 (THO3B) and one control (HO193B) cell line, from day 5 to 14 (**Figures 3G**, **3H and S4G**). The presence of ZO1 at the outer surfaces bending or connecting to internal apical surfaces in both control (day 9) and TS21 (days 5 and 7) cerebellar organoids, suggest that TS21 retains the ability to internalise the outer apical domains (**Figure 3G**). This apical internalisation is consistent with the reduction of outer apical surface percentage, starting from day 5 in TS21 organoids and day 9 in the control (**Figure 3I**). To confirm that the reduction in apical surface and lumen occurs over time, linear regression analysis was performed. While no significant correlation between differentiation time and outer apical surface was observed in the controls, TS21 organoids showed a significant negative correlation, confirming a progressive reduction in outer apical surface across differentiation (**Figure 3I**). In contrast, the inner apical surface significantly decreased over the course of differentiation in both control and TS21-derived organoids (**Figure 3J**). Linear regression analysis also showed that the lumen volume increased significantly over time in the control organoids (**Figure 3K**). No significant correlation was detected in TS21 organoids, suggesting that TS21 organoids fail to undergo normal lumen expansion during differentiation. Together, these results suggest that TS21-derived cerebellar organoids undergo polarisation and lumen morphogenesis but fail to maintain and expand the apical structures during organoid differentiation.

### 4. TS21-derived cerebellar organoids exhibit ECM defects

As ECM regulates the maintenance of neuroepithelial cell polarity, we investigated whether apical defects in TS21-derived organoids are associated with abnormal ECM. Cryosection immunostaining for the main components of ECM (fibronectin, pan-laminin and collagen IV) was performed at day 14 of differentiation. Control organoids started basally accumulating ECM, showing an organised laminal pattern (**Figures 4A**, **4B and S5A**). However, in TS21 condition a disorganised accumulation was found with a diffuse globular-like pattern (**Figures 4A**, **4B and S5A**). Analysis of significantly up-regulated genes in TS21 organoids revealed significantly altered expression of ECM-related genes (**Figure S5B**), as indicated by the enrichment of the collagen-containing ECM Gene ontology term (GO:0062023). As different ECM components are present in the basement membrane and interstitial matrix, and their structure and biochemical composition differentially regulate epithelial function ^29^, their expression was analysed separately. At day 14, TS21-derived organoids showed a significant reduction in transcripts associated with basement membrane, with no significant change in the Z-score of interstitial matrix-related genes (**Figures 4C and S5C**). Nevertheless, *LAMA2, A5, B2, C2*, and *C3*, *COL4A6*, and *ITGA6, ITGB4, ITGA3* were expressed at higher level in the control organoids (**Figure 4D**), suggesting a laminin-332 enrichment and potential α6β4-mediated anchoring. Instead, TS21 organoids exhibited increased expression for *LAMA4*, and *B1, COL4A1-2*, and *ITGB1* (**Figure 4D**), which is indicative of an enriched β1 integrin-associated basement membrane. This data suggests that in TS21-derived cerebellar organoids there is an altered ECM expression and epithelial-specialised basement membrane composition, which may impact downstream signalling and associated tissue polarity and mechanical stability.

**Figure 4.**
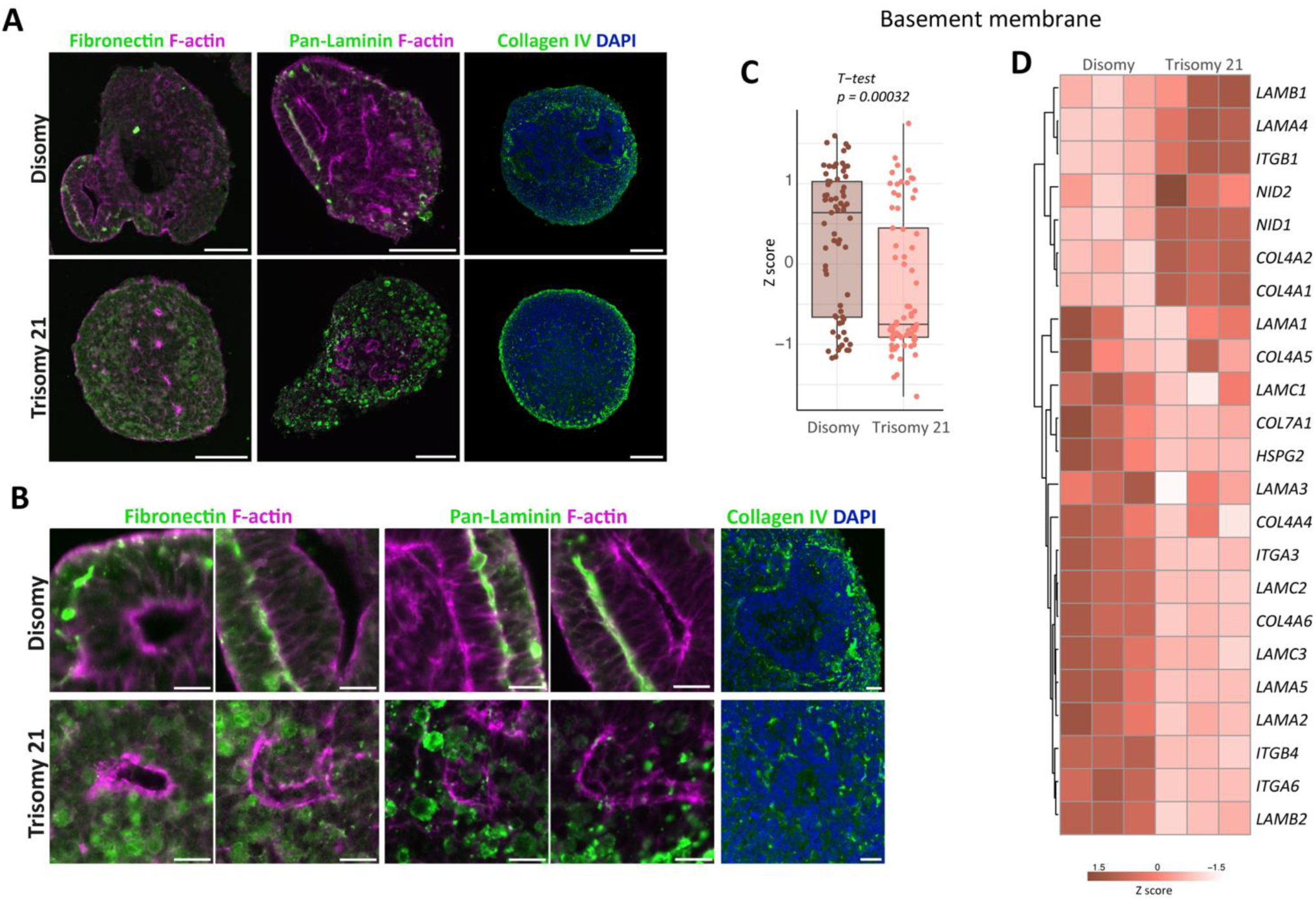
Extracellular matrix (ECM) defects in TS21 cerebellar organoids. **A.** Representative immunofluorescence images of disomy and TS21 showing the distribution of different ECM components, including fibronectin, pan-laminin and collagen IV. Scale bar, 100 μm. **B.** Magnification of ECM components immunostaining showing the ECM location surrounding the neural rosettes. Scale bar, 20 μm. **C.** Box-blot showing the distribution of basement membrane genes (listed in D) obtained from bulk RNAseq of disomy and TS21 organoids at day 14. Data represents minimum to maximum Z-score. **D.** Heatmap of representative basement membrane-related genes from bulk RNAseq data. Values are shown as Z-score.

### 5. Reduced RAB11 apical accumulation in TS21-derived cerebellar organoids

As the maintenance of apical structures and regulation of neural progenitors can be critically dependent on the ECM ^8,30–32^, we next investigated the impact of TS21-derived ECM defects. Focal adhesion kinase (FAK) acts as a primary mechano-sensing mediator activated downstream of integrin-ECM engagement ^33,34^, so we investigated whether the altered ECM organisation in TS21 organoids was associated with changes in FAK signalling. We quantified the proportion of phosphorylated FAK (pFAK, activated ^35^) normalised to the total FAK expression and observed an increased pFAK/total FAK ratio in TS21-derived cerebellar organoids compared with controls (**Figures 5A**, **5B, S6A and S6B**). As pFAK/total FAK ratio alterations can feed into altered down-stream FAK signalling, we next analysed the expression of downstream ECM dynamic-related transcripts that could influence apical stability (**Figure 5C**), including genes involved in cytoskeletal components, Rho GTPases, focal adhesion, and mechano-transduction pathways. While most of these genes were more expressed in TS21 organoids, such as *ACTB, RHOA, ROCK1-2*, and *YAP1*, disomy organoids expressed higher RNA levels for *PTK2*, *PXN* and *VIM* (**Figure 5C**). Together with ECM alterations, the upregulation of transcripts related to ROCK/RHOA signalling could feed into mechano-transduction pathways that regulate cell-matrix adhesion, apical junction organisation, and actomyosin contractility. Indeed, transcriptomic analysis of day 14 TS21 organoids showed a significant upregulation of mRNA expression of actin-related and downregulation of myosin-related genes (**Figure 5D**). Increased expression of actin polymerisation and adhesion-association genes included *PFN1, ARPC1B, VCL* and *TLN1* whilst a decreased expression of myosin V (MyoV) motor genes (*MYO5B* and *MYO5C)* was observed (**Figure 5D**). Immunostaining of organoid cryosections at day 14 was performed and showed F-actin in the ZO1^+^ apical junctions in both disomic and TS21 organoids (**Figure 5E**), consistent with actin enrichment at apical domains. Outside the apical lumens, F-actin labelled cell boundaries across the organoid tissue (**Figure 5E**), and it was found that TS21 organoids also accumulated F-actin in non-apical loci (red arrows). MyoV was broadly detected throughout the control organoids (**Figure S6C**), and higher-magnification views showed altered MyoV organisation at the apical surface in TS21 organoids, where MyoV signal appeared less accumulated relative to disomy (**Figure 5F**). Together, these observations suggest that TS21-organoids display altered actomyosin-associated apical organisation. MyoV transports the RAB11 recycling endosomes along actin filaments ^36,37^, which mediates the return of internalised cadherins to plasm membrane, an essential process for adherens junction maintenance ^38^. We therefore determined whether RAB11 localisation was altered in TS21-derived cerebellar organoids. Immunofluorescence staining for RAB11, ZO1 and F-actin showed that RAB11^+^ endosomes were enriched at the apical surface in the disomy organoids, but not in the TS21 (**Figures 5G**, **5H and 5I**). Quantitative analysis revealed a significant reduction in the proportion of lumens that has RAB11 endosomes in TS21-derived organoids compared with disomic controls (**Figures 5I**, **5J**). The lack of RAB11 endosomes at the apical surface can impair apical proteins turnover and contribute to defective apical junction maintenance in TS21 cerebellar organoids.

**Figure 5.**
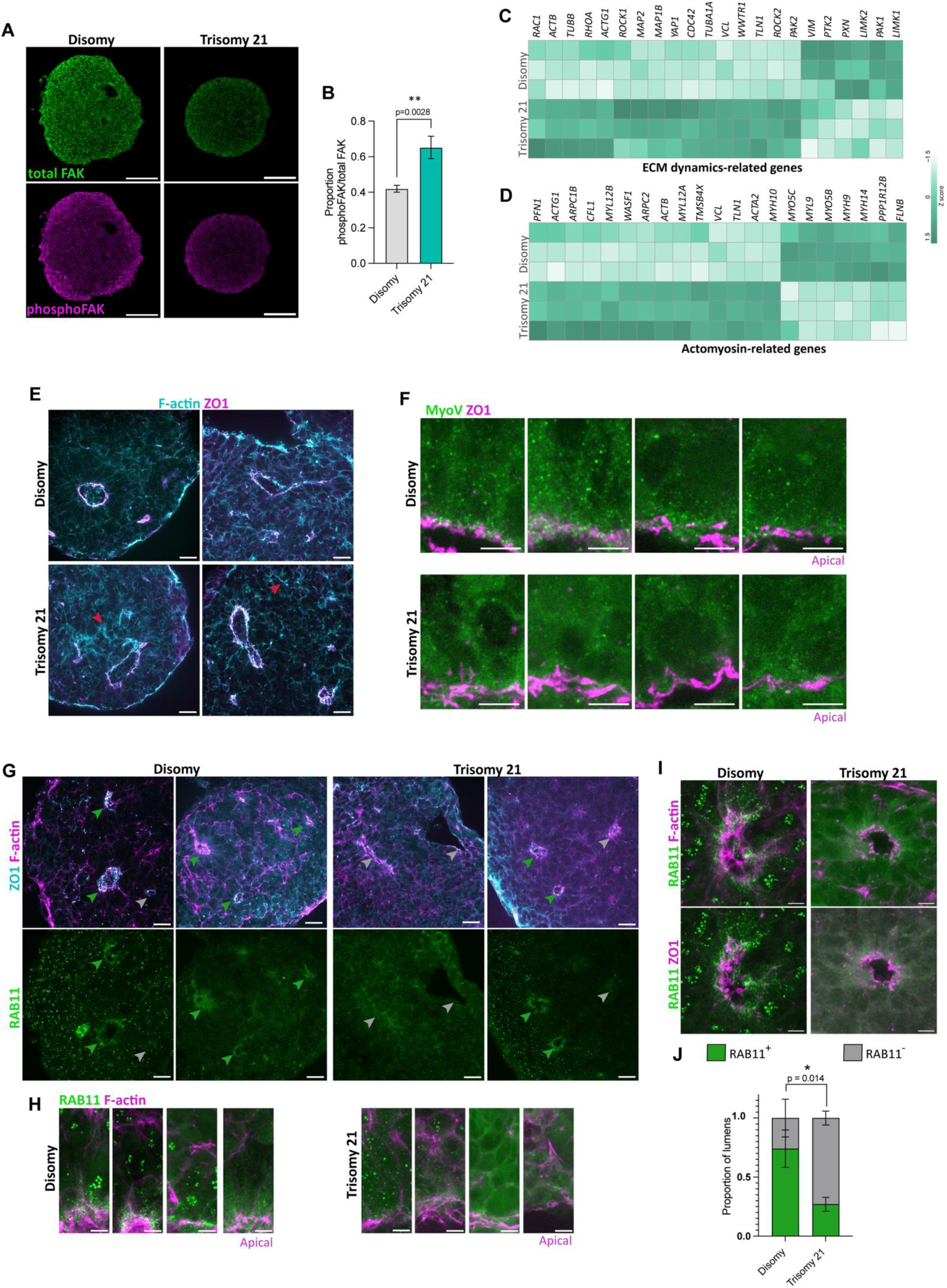
RAB11 defects in TS21 organoids at day 14. **A.** Immunofluorescence staining for total FAK and phosphoFAK in cerebellar organoid cryosections. Scale bar, 100 μm. **B.** Bar-plot showing the ratio of phosphoFAK/totalFAK. Data obtained from 3 disomic and 3 TS21 cell lines. Mann-Whitney test; error bars represent SEM. **p = 0.028. **C.** Heatmap highlighting selected ECM dynamics, including genes that respond to mechanical signals, maintain cytoskeletal structure, or regulate focal adhesion. Values were obtained from bulk RNAseq and are shown as Z-score. **D.** Heatmap highlighting selected actomyosin-related genes. Values were obtained from bulk RNAseq and are shown as Z-score. **E.** Representative staining of organoid cryosections for F-actin and ZO1. Scale bar, 20 μm. The red arrowheads indicate F-actin accumulation in the absence of ZO1 expression in TS21. **F.** Representative high magnification of apical surfaces stained for F-actin and MyoV. Scale bar, 5 μm. **G.** Immunostaining against ZO1, F-actin and RAB11 in cerebellar organoid cryosections. The green arrowheads indicate apical lumens with RAB11 accumulation and grey indicates RAB11-negative lumens. Scale bar, 20 μm. **H.** Representative high magnification images of apical surfaces stained for F-actin and RAB11. Scale bar, 5 μm. **I.** Immunofluorescence staining of organoid cryosections for F-actin, ZO1 and RAB11. Representative apical lumens are shown with RAB11 accumulation at the apical surface or without detectable RAB11 enrichment. Scale bar, 5 μm. **J.** Stacked bar graph shows the proportion of lumen accumulating RAB11. Data obtained from 3 disomic and 3 TS21 cell lines. Two-way ANOVA (Uncorrected Fisher’s LSD); error bars represent SEM. *p = 0.014.

### 6. Matrigel restores apical junction maintenance and progenitor balance in TS21-derived organoids

TS21-derived cerebellar organoids showed apical defects that we hypothesise to arise from a combination of defective ECM signalling and altered RAB11 localisation. Next, to determine whether the apical defects were driven by ECM alterations associated with HSA21 triplication, we tested whether ECM supplementation with Matrigel (MG) treatment could restore apical organisation. For that, organoids were differentiated from day 2 to 14 in a MG-enriched media, specifically 2% MG-enriched media for days 2 and 5, followed by 1% supplementation at day 7. At day 14, after MG treatment, TS21 and disomic organoids showed large ZO1^+^ lumens (**Figure 6A**). Quantification of the five largest lumen volumes per organoid showed a significant lumen expansion following treatment in both conditions (**Figure 6B**). When normalised to total organoid volume, no significant differences were found between disomy and MG-treated disomy (**Figure 6C**), potentially due to the increased organoid size in MG-enriched condition. MG-enriched cultures rescued lumen volume in TS21, with no significant changes detectable between TS21 and disomic controls (**Figures 6B and 6C**). Moreover, MG treatment rescued apical accumulation of RAB11 endosomes in TS21 organoids (**Figure 6D**).

**Figure 6.**
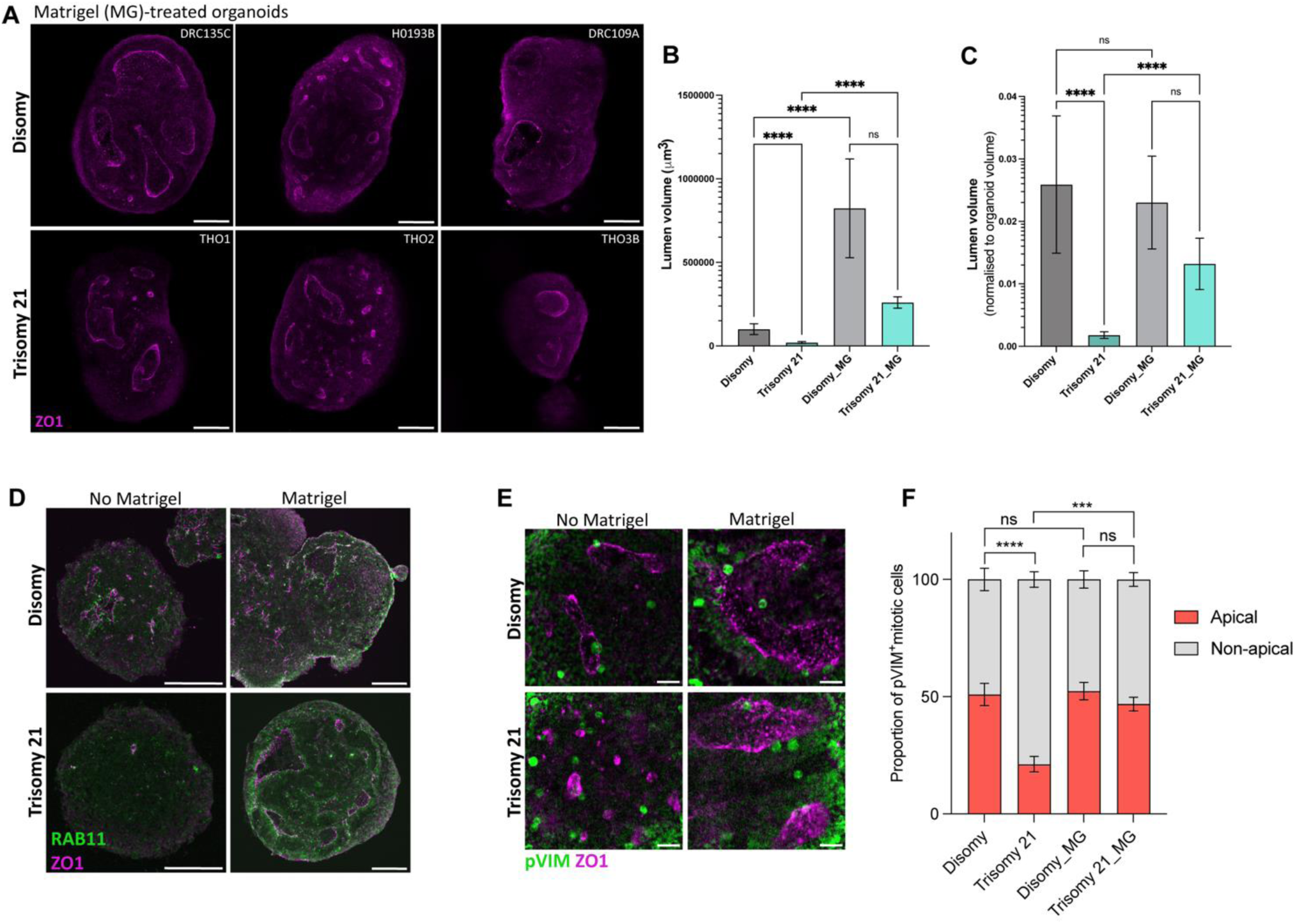
Matrigel restores apical maintenance in TS21 organoids at day 14. **A.** Representative single stack from whole mount immunofluorescence staining for ZO1, showing apical lumens at day 14 after Matrigel treatment. Scale bar, 100 μm. **B.** Bar-plot graph shows lumen volumes at day 14. Data obtained from 3 disomic and 3 TS21 cell lines. Mann-Whitney test; error bars represent SEM. *** p < 0.001, ****p < 0.0001, ns - not significant. MG; Matrigel-treated organoids. **C.** Bar-plot depicts lumen volume normalised to total organoid volume at day 14. Data obtained from 3 disomic and 3 TS21 cell lines. Mann-Whitney test; error bars represent SEM. MG; Matrigel-treated organoids. **D.** Representative staining of organoid cryosections for ZO1 and RAB11. Scale bar, 100 μm. **E.** Representative single stack from whole mount immunofluorescence staining for ZO1 and pVIM at day 14. Scale bar, 20 μm. **F.** Stacked bar diagrams depict the proportion of pVIM^+^ cells proliferating near to apical or non-apical regions at day 14. Data obtained from 3 disomic and 3 TS21 cell lines. Two-way ANOVA (Uncorrected Fisher’s LSD); error bars represent SEM. *** p < 0.001, ****p < 0.0001, ns - not significant.

To determine whether the apical versus non-apical proliferation balance is also restored in MG-treated TS21 cultures, the proportion of apical and non-apical mitotic cells was quantified (**Figure 6E**). MG-treated TS21 organoids showed a significant increase in the proportion of apical mitoses when compared with non-treated TS21 organoids, while disomic and MG-treated disomic organoids had similar proportions of apical and non-apical progenitors (**Figure 6F**). These results suggest that MG treatment restores the apical versus non-apical proliferation balance observed in TS21 organoids, without affecting disomic controls. Together, this data indicates that supplementation with exogenous ECM components is sufficient to ameliorate TS21-associated apical and progenitor type defects, supporting the hypothesis that ECM alterations impair the apical organisation that is observed in TS21 organoids.

### 7. ECM alteration and mechanical defects in TS21 iPSC-derived neuroepithelial sheets

Our results suggest that ECM has a significant impact on apical-basal polarity in TS21-derived cerebellar organoids. We additionally investigated the impact of TS21 on epithelial mechanical properties in a homogeneous 2D system, without confounding effects of overall organoid geometry. To achieve this, three TS21 and four control iPSC lines growing on a MG-coated surface were also differentiated into 2D pseudostratified neuroepithelial sheets, using dual-SMAD inhibition protocol ^39–41^. This simplified 2D neuroepithelial system incorporates an initial exogenous basement membrane component. As previously described and characterised ^24^, after 8 days of differentiation, disomy and TS21 iPSC-derived neuroepithelial cells displayed an organised apical-basal polarity, characterised by apical enrichment of F-actin and formation of a single continuous laminin-containing basement membrane (**Figure 7A**). However, a heterogeneous neuroepithelium was found in TS21, with some regions containing ectopic accumulation of laminin, suggesting a disrupted basement membrane organisation (**Figure 7A**). Sum Z-projection analysis of pan-laminin staining revealed a homogeneous ECM network in the disomy, with relatively even distribution of signal, whereas TS21-derived neuroepithelial sheets showed a heterogeneous pattern of pan-laminin, with areas similar to the control (right **Figure 7B**) and other areas with abnormal pan-laminin distribution (left; **Figure 7B**).

**Figure 7.**
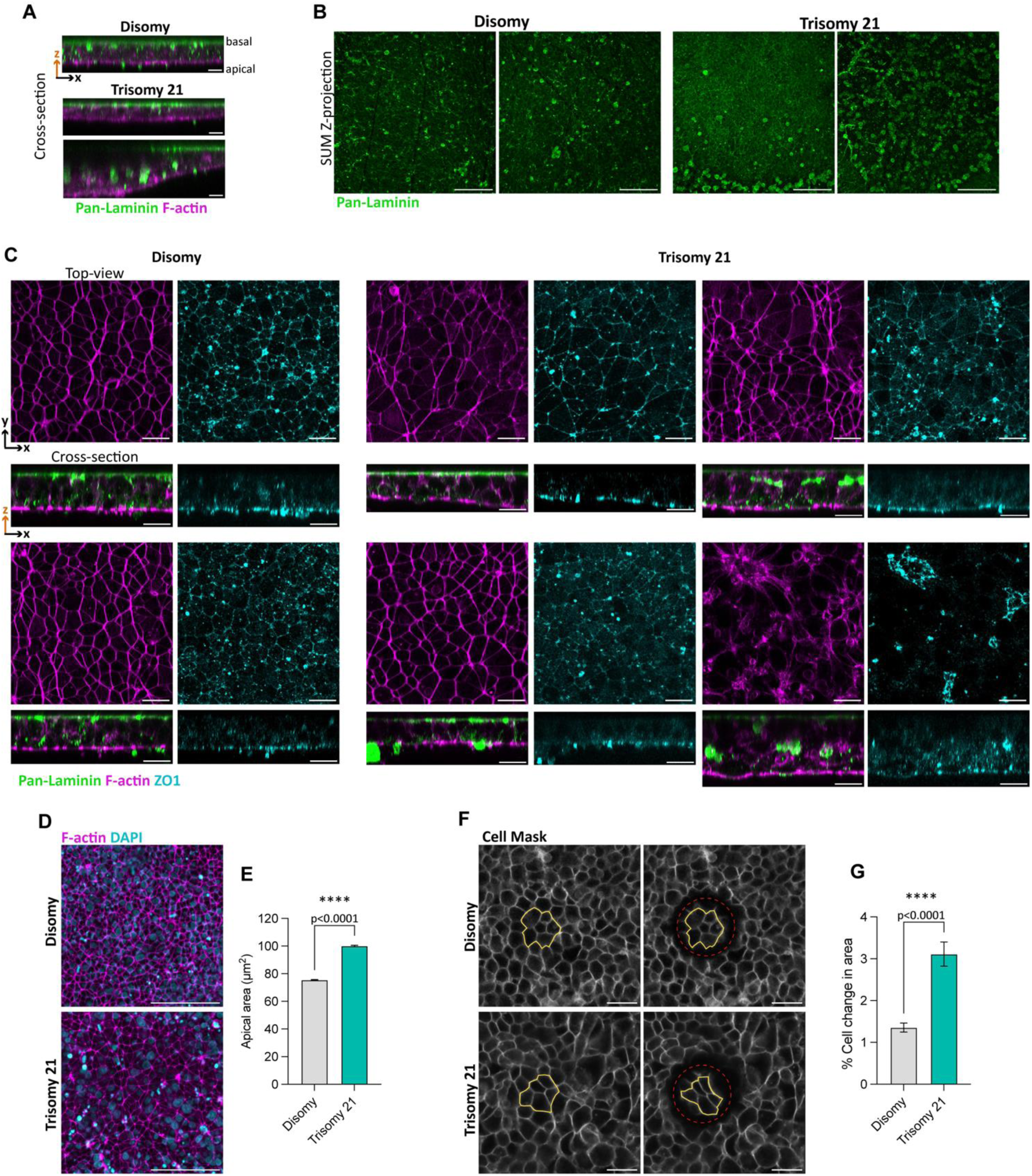
TS21 iPSC-derived neuroepithelial cells exhibits apical and ECM alterations at day 8. **A**. Representative cross-section images of neuroepithelium derived from disomy and TS21, immunostained for pan-laminin and F-actin. Scale bar, 20 μm. **B.** Representative SUM projection images of immunostaining for pan-laminin. Scale bar, 20 μm. **C.** Representative top-view and cross-section images of day 8 cultures from disomy and TS21, stained against pan-laminin, F-actin and ZO1. Scale bar, 20 μm. **D.** Staining for F-actin showing cell apical surfaces at day 8. Scale bar, 100 μm. **E.** Bar-plot depicts the mean cell apical surface area in disomy and TS21 neuroepithelia. Data obtained from 4 disomic and 3 TS21 cell lines. Mann-Whitney test; error bars represent SEM. ****p < 0.0001. **F.** Representative live-imaging of CellMask-stained day 8 neuroepithelial cultures before and after a linear circular laser ablation, indicated by red dash line. Yellow outlines the cell cluster before and after the ablation. **E.** Bar-plot diagram shows the % cell change area. Data obtained from 4 disomic and 3 TS21 cell lines. Mann-Whitney test; error bars represent SEM. ****p < 0.0001.

To determine whether apical junctions were also affected, we additionally stained day 8 neuroepithelial sheets with ZO1 (**Figure 7C**). In orthogonal views, disomic cultures exhibited clear apical-basal organisation, with pan-laminin positioned basally and F-actin/ZO1 enriched at the opposite apical surface (**Figure 7C**). In contrast, TS21-derived neuroepithelium showed more variable and altered apical-basal polarity. In some regions ZO1 apical accumulation was reduced or disrupted, coinciding with ectopic pan-laminin accumulation (**Figure 7C**). Unlike cerebellar organoids, iPSC-derived pseudostratified neuroepithelial sheets have apical surfaces that are accessible for imaging and quantification ^24^. Thus, F-actin staining was used to label the apical junctional end-feet of neuroepithelial cells and to select the regions with preserved epithelial cytoarchitecture for analysis (**Figures 7C**, **7D**). Quantitative analysis of apical surface area of individual progenitor cells demonstrated that TS21-derived neuroepithelia exhibited significantly larger apical surfaces compared to controls (**Figure 7E**). To determine whether this increase in cell apical surface area reflected alteration on neuroepithelial tissue mechanics, circular laser ablation (**Figure 7F**) and recoil analysis were performed as previously described ^24,42^. Recoil is defined by the change of individual cell apical area after laser ablation as a readout of mechanical tension. TS21-derived neuroepithelial sheets showed significantly higher apical recoil than controls, demonstrated by the reduction in cell area (**Figure 7G**), suggesting altered apical mechanical properties. Together, these findings suggest that TS21-derived neuroepithelial sheets exhibit intrinsic mechanical abnormalities. The TS21-associated apical phenotype appeared milder than the apical defects observed in cerebellar organoids, suggesting that the initial MG-coating used for iPSC attachment may support the apical-basal maintenance.

### 8. Apical alterations in the human DS foetal cerebellum

To determine whether ECM, apical surface and RAB11 were also disrupted in human TS21 cerebellum, foetal tissue at 13 and 17 post-conceptional weeks (PCW) were analysed. Immunostaining for pan-laminin and collagen IV showed the presence of ECM components in blood vessels at 13 and 17 PCW (**Figure 8A**). Gross apical junctional architecture in the TS21 cerebellum was largely intact, with apical proteins, ZO1 and NCAD, and F-actin expressed at the apical surface (**Figures 8A and 8B**). High resolution imaging showed NCAD and RAB11 containing vesicles surrounding and docking at the F-actin labelled apical junctions in the control foetus (**Figure 8C**). This pattern of distribution of NCAD and RAB11 was less evident in TS21 cerebellum (**Figure 8C**), suggesting that either trafficking or levels of expression are down regulated. These findings suggest that, in contrast to cerebellar organoids, apical surface alterations are less severe in DS foetal brains. The potential interaction of apical pVIM⁺ mitotic progenitors with blood vessels (**Figure S7**) suggests that local vascular-associated ECM may provide additional structural support, which can contribute to maintain the gross apical junctional architecture observed in the developing human cerebellum.

**Figure 8.**
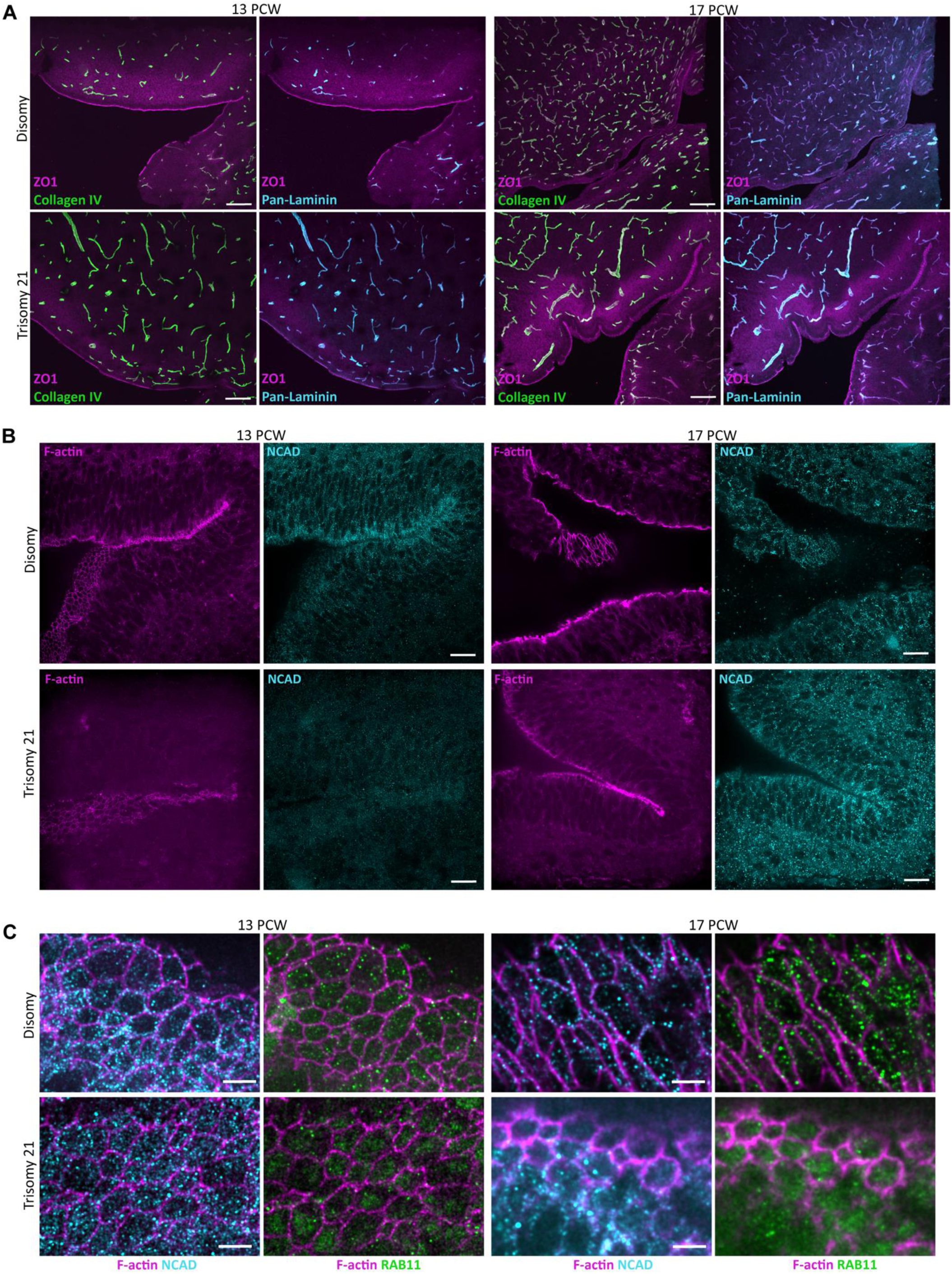
Characterisation of human foetal DS cerebellum (13-17 PCW). **A.** Immunofluorescence staining for ZO1, collagen IV and pan-laminin in human cerebellum cryosections at 13 and 17 PCW. Scale bar, 200μm. **B.** Representative staining for F-actin and N-cadherin (NCAD) in human cerebellum cryosections at 13 and 17 PCW. Scale bar, 20μm. **C.** Immunofluorescence images for NCAD, F-actin and RAB11 from disomy and TS21at 13 and 17 PCW. Scale bar, 5μm.

## Discussion

This study provides insight into how triplication of HSA21 impacts human brain development. This has been achieved using human iPSC-derived cerebellar organoids, 2D pseudostratified neuroepithelial models, and human foetal cerebellar tissues. We have consequently been able to uncover the mechanism underlying the TS21-derived apical disruption and ECM alterations previously reported iPSC-derived neural model and human developing neocortex ^12,22^. Here, we demonstrate that TS21 disrupts RAB11 apical accumulation, leading to defects in apical maintenance and progenitor type balance. Our human iPSC-derived neural models reveal that alteration of ECM composition and spatial distribution can perturb down-stream signalling pathways important for RAB11^+^ vesicle trafficking along actin filaments.

RAB11^+^ recycling endosomes are known to regulate trafficking and localisation of apical polarity proteins, as well as the maintenance of adherens junctions, across several model systems ^38,43,44^. The disruption of RAB11 localisation or function compromises apical junction stability ^38,45,46^, leading to loss of apical junctions integrity and premature neuronal differentiation ^47^. Our TS21 organoids retain key apical junction proteins on existing apical surfaces, including ZO1, PAR6 and NCAD. This indicates that apical identity is not completely abolished. Instead, RAB11 mis-localisation may impair the trafficking, recycling or turnover of specific cargoes required for maintenance of the apical surface.

Our study suggests a potential ECM-focal adhesion-trafficking mechanism underlying apical changes in TS21-derived organoids. Transcriptomic analysis of TS21 cerebellar organoids further reveals altered expression of transcripts associated with ECM dynamics, including cytoskeletal components, Rho GTPases, and focal adhesion pathways in the TS21 condition. Together with ECM component mis-localisation, increased expression of transcripts related with ROCK/RHOA signalling ^48^ suggests altered cytoskeleton tension, which may feed into altered FAK signalling activation. This is supported by our data showing that TS21 cerebellar organoids have an increased phosphoFAK/total FAK ratio when compared to controls. Activated FAK can promote the activation of paxillin ^33,49,50^, a key focal adhesion scaffold protein involved in cytoskeletal remodelling downstream to ECM-cell interactions ^33,51,52^. Paxillin regulates actin cytoskeleton organisation through its interaction with actin-binding proteins and by modulating members of the Rho GTPase family ^53^. TS21-derived organoids show a significant altered expression of actin- and myosin-related genes, including downregulation of genes encoding class V myosin motors, accompanied by reduced apical MyoV localisation. This is relevant because recycling endosome transport requires coordinated activity between Rab small GTPases and class V myosin motors ^54,55^, including the transport of RAB11^+^ recycling endosomes along actin filaments ^36,37^. Notably, paxillin, a downstream mediator of FAK signalling, has previously been shown to regulate the RAB11/MyoV-dependent trafficking machinery involved in apical-basal polarity in mammary gland morphogenesis ^56^. Our data supports this model by showing a loss of RAB11 and MyoV at the apical surface in TS21, suggesting disruption of apical trafficking. One potential hypothesis is that ECM-associated alterations in TS21 organoids affect FAK-paxillin signalling, actomyosin organisation and RAB11/MyoV-dependent trafficking. This will then contribute to impaired apical polarity maintenance. Further work will be required to identify the affected RAB11-dependent cargoes and to determine whether RAB11 mis-localisation is a downstream consequence of altered trafficking.

The apical instability observed in our TS21 organoids was associated with a reduction in the proportion of apical progenitors and a relative increase in non-apical progenitor populations. Other studies have previously demonstrated that ECM-integrin signalling can regulate neural progenitor maintenance ^57^, including apical radial glia cell attachment to the ventricular ^58^ and basement membrane surfaces ^7^, asymmetric division ^58^, and intermediate progenitor cell proliferation ^32^. This suggests that ECM defects can visibly impact the expansion of progenitor populations. In addition, the RAB11-associated apical instability observed in TS21 raises the possibility that disruption of RAB11-mediated apical integrity may contribute to altered progenitor balance. Impaired apical localisation of RAB11 in TS21-derived organoids can lead to apical junction instability, promoting the detachment of apical progenitors and their subsequent transition into basal progenitors and neurons ^47^. This process could ultimately affect tissue growth and contribute to reduced cerebellum volume, a characteristic feature of DS.

Our ECM supplementation experiments support a functional role for ECM defects in the TS21 phenotype. Exposure of cortical organoids to extrinsic ECM components, such as MG, has been shown to modulate tissue morphogenesis ^59^ by promoting cell polarisation and supporting lumen maintenance ^60^. In our model, MG supplementation restored apical junction organisation in TS21-derived organoids and rescued associated proliferative and progenitor phenotypes, without affecting the control. This selective rescue supports the interpretation that ECM defects are functionally involved in driving apical instability in TS21 organoids.

Analysis of DS foetal cerebellum revealed a less severe apical phenotype, suggesting partial buffering of apical defects *in vivo*. In contrast to TS21 cerebellar organoids, the developing DS cerebellum showed largely preserved gross apical organisation. Previous study reported truncated radial glia, containing a short basal process ending onto a laminin-expressing blood vessels in newborn ferret cerebral cortex^61^. Then, one plausible explanation for subtle phenotype in DS foetal cerebellum is that blood vessels provide local ECM support ^62,63^ that helps stabilise epithelial organisation during development. These structures are largely absent from current organoid models, which may make ECM-sensitive phenotypes more apparent *in vitro*. Although gross epithelial architecture was preserved, changes at the cellular level were observed in developing DS cerebellar tissue. Whereas control foetal cerebellum showed predominantly small punctate RAB11^+^ structures, consistent with recycling endosomes and transport vesicles ^64,65^, TS21 foetal cerebellum showed less punctated RAB11^+^ distribution. This altered distribution may indicate changes in apical membrane trafficking, junctional remodelling, or vesicle recycling dynamics in TS21. In parallel, NCAD appeared less clearly enriched or docked at apical junctions in TS21 cerebellar tissue, suggesting a potential impairment of NCAD apical trafficking. Although apical architecture is not overtly disrupted in the TS21 foetus, progenitors may display altered apical membrane trafficking and junctional organisation. This supports the view that *in vivo* components, such as blood vessels, may compensate for, but not fully eliminate, intrinsic defects in apical domain organisation. Furthermore, significant alterations in vascular ECM gene expression have been reported in the developing human DS neocortex ^12^, suggesting that although vascular ECM components are present, their altered composition may prevent them from fully compensating for the neuroepithelial defects.

Our 2D pseudostratified neuroepithelial cultures provide an important bridge between the organoid and foetal tissue phenotypes. In these cultures, the initial MG coating may provide extracellular cues that support apical-basal polarity and adherens junctions and partially rescue severe epithelial disorganisation. Although overall epithelial organisation was maintained, ECM mis-localisation, altered apical surface organisation, and abnormal mechanical properties were still observed in some regions of the TS21-derived neuroepithelium.

Altogether, our findings suggest that extrinsic ECM can buffer severe epithelial disorganisation, while intrinsic TS21-associated defects in ECM organisation, apical trafficking and biomechanical regulation still persist. Our results highlight the value of the iPSC-derived neural models as platforms for mechanistic and functional interrogation. Especially, these systems have revealed ECM defects and disrupted apical organisation in TS21-derived neural cultures. These findings subsequently guided the analysis of apical defects and RAB11-associated trafficking in human foetal tissues, supporting the relevance of phenotypes observed *in vitro*. While our experimental models are inherently reductionist, they can provide a powerful experimental framework to uncover disease-associated cellular defects and to investigate their potential relevance during human neurodevelopment.

In conclusion, our data predicts that TS21 disrupts the coordination between ECM organisation, apical junction maintenance and mechanics during early neural development. Rather than causing a complete failure of epithelial architecture, TS21 appears to produce an instability of the apical maintenance, which becomes particularly evident in reductionist *in vitro* systems that lack the full complexity of the embryonic microenvironment. The rescue of apical and progenitor phenotypes by MG further supports the view that ECM-dependent signalling and structural support are important modifiers of TS21 neural phenotypes. Together, these findings propose that impaired ECM-progenitor niche interactions represent a previously underappreciated RAB11-dependent mechanism that underlies to altered neuroepithelial development in DS.

### Limitations of the study

Although our data indicate altered mechanics in TS21 neural tissue, we have not yet determined whether tissue stiffness and apical tension are directly affected, nor whether similar biomechanical changes occur in human foetal tissue. The limited number of human foetal samples analysed restricts the extent to which these observations can be generalised. Direct biomechanical measurements in iPSC-derived models and, where feasible, in larger cohorts of foetal samples will be required to resolve these questions. The limited availability of human foetal tissue also means that subtle apical phenotypes observed *in vivo* should be interpreted as supportive rather than definitive evidence of altered apical regulation in DS cerebellar development.

In addition, although ECM supplementation rescues key phenotypes, this does not establish ECM dysregulation as the initiating event in TS21 neural phenotype. Other cell-intrinsic effects of HSA21 triplication are likely to contribute and may interact with ECM-dependent pathways. Finally, iPSC-based models cannot fully recapitulate the complexity of the *in vivo* environment, however they provide powerful access to early human neurodevelopmental processes and are useful to reveal disease mechanisms.

## Acknowledgements

This study and T.P.S. were supported by Great Ormond Street Hospital Children’s Charity Michael Uren Foundation (project reference V4521) and Child Health Research CIO (20-21/SI5). Wellcome Human Developmental Biology Initiative (215116/Z/18/Z) supported the HDBI imaging Hub and the experimental work on the human tissue. I.S. and G.C. are both funded by NIHR BRC GOSH. ZO1-EGFP iPSC line purchased with RS grant RG\R2\232082. We would like to acknowledge Human Developmental Biology Resource (HDBR) for providing human samples. UCL and NIHR Great Ormond Street Hospital Biomedical Research Centre for supporting facilities at UCL GOS ICH. A special thanks goes to Dale Moulding that provided support and advice on imaging acquisition, and Alexandre Baffet lab, namely Ryszard Wimmer and Clarisse Brunet Avalos, for teaching T.P.S. to perform organotypic slice cultures of organoids.

## Contributions

T.P.S. and P.A. conceived the project, supervised the work, and wrote the manuscript. P.H. and N.D.E.G. helped with the project design. F.J.L. and J.S. contributed with SFC086-03-01, DRC109A, DRC135C, THO1, THO2, THO3B iPSC lines. G.G.G. and P.D.C. contributed with H0193B iPSC line. G.G. contributed with mEGFP-ZO1 iPSC line. J.C., G.G. and N.D.E.G. contributed with reagents. T.P.S. performed the experiments with iPSC-derived cerebellar organoids and most of the respective analysis. E.R.I. performed human tissue staining and analysis. G.G. developed script in Fiji to separate inner outer layer in organoids and provided input on the biomechanics. E.U., D.H. prepared and processed human samples. T.P.S and I.A. contributed to 2D culture, laser ablation and analysis. G.C. and I.S. helped with the immunostaining of organoids. All authors commented on the manuscript.

## Methods

### Human induced pluripotent stem cell culture

In this study, five distinct control iPSC lines (H0193B ^24,25,66^, SFC086-03-01 ^24,25^, DRC109A, DRC135C ^18,67^, mEGFP-ZO1) and three TS21-derived iPSC lines (THO1, THO2, THO3B) ^18,67^ were used. A summary of detailed information about human iPSCs lines used on this study can be found in **Supplementary Table S1**. All cell lines were cultured on Matrigel (Corning)-coated 6-well plates with mTeSR™ Plus medium (StemCell Technologies) at 37°C in 5% CO2. Cells were passaged when the colonies covered approximately 85% of the surface area of the culture dish, using EDTA in DPBS (final concentration 0.5 mM, Thermo Fisher Scientific). All cell lines were maintained below passage 40 and periodically tested for mycoplasma contamination and copy number variation.

### Cerebellar organoid differentiation

Cerebellar organoids were generated using a previously described protocol ^23^. Briefly, for single-cell seeding, cells were treated with accutase (Sigma) for 5 min at 37°C. After dissociation, single-cells were seeded in Aggrewell 800 plates (StemCell Technologies) at a density of 1500 cells/aggregate in mTeSR™ plus medium (StemCell Technologies) supplemented with Y-27632 (10 μM, StemCell Technologies). After 24 hours, medium was replaced and aggregates were maintained in mTeSR™ plus without Y-27632 for another 24 hours. From day 2 to day 21 after seeding, gfCDM was used as basal medium for the differentiation ^23^. Recombinant human basic FGF (FGF2, 50 ng/ml, Thermo Fisher Scientific) and SB431542 (10 μM, Sigma) were added to culture on day 2. Full-volume medium replacement with gfCDM (supplemented with FGF2 and SB431542) was performed on day 4. On day 7, medium was fully replaced, supplemented with 33.4 ng/mL FGF2 and 6.7 μM SB431542. Recombinant human FGF19 (100 ng/mL, Thermo Fisher Scientific) was added to culture on day 14 post-seeding, and full-volume replacement was performed on day 18. From day 21, the aggregates were cultured in Neurobasal medium (Thermo Fisher Scientific) supplemented with GlutaMax I (Thermo Fisher Scientific), N2 supplement (Thermo Fisher Scientific), and 50U/ml penicillin/50μg/ml streptomycin (PS, Thermo Fisher Scientific). Full-volume replacement was performed every 7 days. Recombinant human SDF1 (300 ng/ml, Thermo Fisher Scientific) was added to culture on day 28. Matrigel treatment was performed using a previously described modified protocol ^60^. For that, gfCDM (supplemented with FGF2 and SB431542) with 2% dissolved Matrigel was supplied on day 2, followed by full-volume replacement with 2% Matrigel (Corning) supplemented media. On day 7, medium was fully replaced with 1% Matrigel.

### Dual SMAD inhibition neural induction

Differentiation of human iPSCs into neuroepithelial sheets was performed using a dual SMAD inhibition protocol, as previously described ^39–41^. Briefly, iPSCs were lifted with 0.5 mM EDTA and plated at approximately 300,000 cells/cm² in mTeSR™ Plus medium. After 24 hours, once 100% confluency was reached, the medium was replaced daily with 2 mL of neural maintenance medium (NMM) supplemented with 1 μM dorsomorphin (Tocris) and 10 μM SB431542 for a total of 8 days. If cells did not reach 100% confluency, the medium was replaced with fresh mTeSR™ Plus for an additional 24 hours, after which differentiation was initiated.

### Live imaging in cerebellar organoids

To follow lumen formation in cerebellar organoids, mEGFP-TJP1-cl20 (mEGFP-ZO1) were differentiated for 14 days and transferred to glass bottom dishes for imaging. Daily time-lapse imaging was performed from day 2 to day 14, with images acquired every 20 min over periods of more than 14 h. For live imaging, organoids were placed under the microscope in the culture medium at 37C and in a humidified gas-controlled chamber (%CO2 /20%O2). Imaging was performed using a spinning disk CREST V3 - ECLIPSE T2i Nikon inverted confocal microscope, fitted with a long-working-distance ×20 Plan Fluor ELWD dry objective (NA 0.45; WD 6.9–8.2 mm; Nikon) and a Photometric Kinetix camera. Videos were processed using NIS-Elements software and Fiji software and maximum Z-projections.

### Infection of cerebellar organoids

Day 16 organoids were embedded in 4% low-melting agarose and then sliced using a Leica VT1000S vibratome (150-μm-thick slices) in ice-cold DMEM-F12. Organoid slices were transfected with 5μL of pGS-LVV-EF1a:EGFP lentivirus (6.27 x 10^7^ particles/mL) in 1mL of culture media for 2h at 37°C. After 2 h of incubation, slices were washed three times with DMEM-F12 and grown in culture medium at 37°C for 48h, on Millicell cell culture inserts (Merck). Lentiviral vectors were generated at the CRICK Institute, London.

### Cryosectioning and immunofluorescence staining of cerebellar organoid and human tissue slices

Organoids were fixed in 4% PFA either overnight at 4°C or at room temperature for 2 hours and washed in PBS (3×10 min). Human samples were fixed in 10% Formalin at 4°C for at least 2 days. For cryostat processing, samples were incubated overnight in 15% (v/v) sucrose in PBS, embedded in gelatine (7.5% v/v gelatine, 15% v/v sucrose), and then frozen at -80°C. Sections with approximately 15-μm in thickness were cut on a cryostat microtome (Leica CM1950, Leica Microsystems), collected on Superfrost^TM^ Microscope Slides (Thermo Scientific), and stored at −20◦C. Finally, sections were de-gelatinized for 45 min in PBS at 37°C before being processed for immunohistochemistry. Organoids and human slices were then incubated in 0.1 M Glycine (Millipore) for 10 min at room temperature (RT), permeabilized with 0.1% Triton X100 (Sigma) for 10 min at RT, and blocked with blocking permeabilizing solution (0.1% Triton X-100 and 6% BSA in PBS) for 2 hours at RT. Next, samples were incubated overnight at 4°C with the primary antibodies (**Supplementary Table S2**) diluted in blocking solution. Secondary antibodies (**Supplementary Table S3**) were incubated for 1 hour at RT. For human tissue secondary antibodies were incubated overnight at 4°C. Nuclei were counterstained with 4′,6-diamidino-2-phenylindole (DAPI, 1.5 μg/mL, Severn Biotech Ltd 17057). For phalloidin staining, sections were incubated with Alexa Fluor™-568 or Alexa Fluor™-647 conjugated phalloidin (1:100 in blocking solution for 1 hour, Thermo Fisher Scientific). Finally, samples were mounted in VECTASHIELD antifade mounting media (2BScientific).

### Whole-mount staining of organoids

Organoids were fixed in 4% PFA either overnight at 4°C or at room temperature for 2 hours and washed in PBS (3×10 min). For whole mount immunofluorescence, a previously described modified protocol was applied ^68,69^. All incubations were performed with gentle agitation unless otherwise stated. Organoids were dehydrated through an increasing methanol series for 1 h each in 25%, 50%, 75% and 100% methanol (diluted in ddH₂O), and left in 100% methanol overnight. Samples were then rehydrated through 75%, 50% and 25% methanol and 1× PBS, for 1 h each, before overnight incubation in 0.5 M acetic acid (Sigma, 64-19-7) at 4°C. The following day, samples were incubated in ECM solution (4 M guanidine hydrochloride – Sigma 50-01-1, 0.05 M sodium acetate – Sigma 127-09-3, and 2% Triton X-100 in 1× PBS) at RT during the day, followed by overnight incubation in 5% CHAPS (in ddH₂O) at RT. Samples were blocked in blocking solution, consisting of PTWH solution (0.2% v/v Tween-20 and 10 mg/mL heparin in 1x PBS) supplemented with 6% donkey serum and 10% DMSO at RT for at least 6 hours. Then incubated with primary antibodies diluted in antibody solution (PTWH solution with 3% donkey serum and 5% DMSO) for 4 days at 4°C. After 4-6 washes in PTWH for 30 min each, samples were incubated with secondary antibodies diluted in antibody solution for 6 days at 4°C, protected from light. Samples were then washed 4-6 times in PTWH. For embedding, organoids were transferred to cryomolds, and samples were embedded in 1.5% (v/v) agarose prepared in ddH₂O. After solidification, agarose blocks were cut into approximately 5 × 5 mm pieces. Samples were cleared using the MACS Clearing Kit. Embedded samples were dehydrated in 50% ethanol/2% Tween-20, 70% ethanol/2% Tween-20 and 100% ethanol/2% Tween-20 under slow rotation. Samples were then transferred individually to polypropylene tubes containing MACS Clearing Solution and incubated at room temperature under rotation until transparent. The clearing solution was replaced with ethyl cinnamate by three 5 min washes, and samples were mounted on single-depression slides in ethyl cinnamate for imaging.

### Immunofluorescence staining of neuroepithelium

Adherent neuroepithelial cultures were fixed for 15 minutes in cold 4% paraformaldehyde (PFA) and followed by washing in Phosphate buffered saline (PBS, 0.1M). Cells were permeabilized with 0.3% Triton X100 (Sigma) for 15 min at RT and blocked blocking permeabilizing solution (0.1% Triton X-100 and 6% BSA in PBS) for 6 hours at RT. Next, the cells were incubated overnight at 4°C in a blocking-permeabilizing solution containing primary antibodies. The next day, the cells were rinsed three times in PBS before incubation with secondary antibody in blocking-permeabilizing solution for 1h at RT. Finally, the secondary antibody solution was washed off with PBS. Nuclei were labelled with DAPI, and F-actin was labelled with Alexa Fluor™-647 conjugated phalloidin.

### Image acquisition and analysis

Fluorescence images were acquired with Zeiss Examiner LSM 880 point-scanning confocal microscope or spinning disk CREST V3 - ECLIPSE T2i Nikon inverted confocal microscope, fitted with a silicon oil Apochromat Lambda S 100x objective (NA 1.35; WD 0.31-0.28 mm) and a Photometric Kinetix camera.

1. **Organoid area analysis:** To monitor changes in organoid size over time in culture, brightfield images were acquired at different time points using an Olympus IX73 microscope equipped with an ORCA-Flash4.0 LT Hamamatsu C11440 digital camera. Organoid area was quantified in Fiji/ImageJ by manually delineating the outline of each organoid using the polygon selection tool, followed by measurement of the enclosed area.
2. **Lumen volume analysis:** images were acquired using a Zeiss Examiner LSM 880 confocal microscope and then Airyscan processing was performed using Zeiss software. The five largest lumens per organoid were manually selected in Fiji/ImageJ. Each lumen was manually delineated across representative z-sections, and the resulting ROIs were propagated across the z-stack to generate a full 3D lumen segmentation. Lumen area was measured in each optical section, and total lumen volume was calculated by integrating the measured areas across the Z-stack using the Z-step interval.
3. **ZO1 apical surface quantification:** images were acquired using a Zeiss Examiner LSM 880 confocal microscope. Airyscan processing was performed using Zeiss software. To quantify ZO1 distribution in inner and outer organoid regions, a Fiji/ImageJ-based segmentation pipeline was used. For each organoid, the ZO1 image stack was duplicated to generate separate images for whole-organoid masking, outer-region segmentation and inner-region segmentation. The organoid outline was first identified by automatic thresholding using the Huang method, followed by conversion to a binary mask. Bright and dark outliers were removed from the mask, and the organoid area was detected using particle analysis across the stack. To separate outer and inner ZO1^+^ regions, the organoid mask was eroded by 15 pixels. In one duplicated stack, the eroded mask was used to clear the internal region, leaving only the outer rim of the organoid. In a second duplicated stack, the same eroded mask was used to clear the region outside the mask, retaining the inner/core region. The resulting outer and inner stacks were then merged as separate channels to allow visualisation and downstream quantification of ZO1 signal in each compartment. ROIs generated from the segmentation were used to determine the organoid area in each optical section, and total organoid volume was calculated by integrating the measured areas across the Z-stack using the Z-step interval. To quantify inner and outer ZO1^+^ regions, background was measured and background-corrected ZO1 signal was thresholded using a log2 fold-change cut-off of 1 over background, followed by conversion to a binary mask. The area of ZO1^+^ signal was measured in each optical section and integrated across the z-stack to calculate the volume of inner and outer ZO1 expression.
4. **3D quantification of apical progenitors:** images were acquired using a Zeiss Examiner LSM 880 confocal microscope. Airyscan processing was performed using Zeiss software. For pVIM^+^ cell quantification, maximum Z-stack projections were generated from blocks of 10 consecutive optical sections. These projection blocks were separated by 10 Z-sections (20-40 μm) throughout the organoid volume to minimise the risk of counting the same cell more than once.
5. **FAK expression analysis**: Images were acquired using a Zeiss Examiner LSM 880 confocal microscope equipped with a 20×/NA 1.0 Plan-Apochromat water objective. Airyscan processing was performed using Zeiss software. In Fiji/ImageJ software, SUM Z-projections were generated and used to measure the mean fluorescence intensity of total FAK and phospho-FAK within the same organoid.
6. **Apical area quantification:** apical area analyses were performed on F-actin-stained iPSC-derived neuroepithelial sheets and ZO1-stained human cryosections. The F-actin or ZO1 channel was segmented using the CellPose Cyto2 model with calibrated segmentation settings, and the resulting masks were saved as ROIs. The area of each ROI was then measured in Fiji/ImageJ. Approximately three images were acquired per plate and treated as technical replicates.
7. **Deconvolution of human F-actin, RAB11 and NCAD staining images:** Images were acquired using a spinning disk CREST V3 - ECLIPSE T2i Nikon inverted confocal microscope, with a silicon oil Apochromat Lambda S 100x objective (NA 1.35; WD 0.31-0.28 mm) and a Photometric Kinetix camera. For imaging deconvolution, Huygens Essential (version 26.04, Scientific Volume Imaging, Hilversum, The Netherlands) was used to remove noise and reassign out-of-focus light with a theoretically calculated point spread function, using the classic maximum likelihood estimation (CMLE) deconvolution algorithm. Processed image stacks were saved in 32-bit OME-TIFF format.

### Expression profiling with RNA sequencing

1. **Sample collection and RNA extraction**: H0193B and THO3B iPSC line-derived organoids were collected on day 14, using 3 technical replicates. For RNA extraction, organoids were mechanically dissociated and homogenised in 800 μL of TRI reagent (Zymo Research) and then stored at -80°C. Total RNA was extracted from samples using Direct-zol^TM^ RNA Miniprep kit (Zymo Research, R2051), according to the manufacturer instructions.
2. **RNA-seq sample preparation and sequencing:** Sample preparation and sequencing was performed by GENEWIZ NGS Europe (Azenta Life Sciences). RNA samples were quantified using Qubit 4.0 Fluorometer (Life Technologies, Carlsbad, CA, USA) and RNA integrity was checked with RNA Kit on Agilent 5300 Fragment Analyzer (Agilent Technologies, Palo Alto, CA, USA). RNA sequencing libraries were prepared using the NEBNext Ultra RNA Library Prep Kit for Illumina following manufacturer’s instructions (NEB, Ipswich, MA, USA). Briefly, mRNAs were first enriched with Oligo(dT) beads. Enriched mRNAs were fragmented according to manufacturer’s instruction. First strand and second strand cDNAs were subsequently synthesized. cDNA fragments were end repaired and adenylated at 3’ends, and universal adapters were ligated to cDNA fragments, followed by index addition and library enrichment by limited-cycle PCR. Sequencing libraries were validated using NGS Kit on the Agilent 5300 Fragment Analyzer (Agilent Technologies, Palo Alto, CA, USA), and quantified by using Qubit 4.0 Fluorometer (Invitrogen, Carlsbad, CA). ERCC RNA Spike-In Mix reagent (Cat: 4456740) from ThermoFisher Scientific, was added to normalized total RNA prior to library preparation following manufacturer’s protocol. The sequencing libraries were multiplexed and loaded on the flow cell on the Illumina NovaSeq 6000 instrument according to manufacturer’s instructions. The samples were sequenced using a 2×150 Pair-End (PE) configuration v1.5. Image analysis and base calling were conducted by NovaSeq Control Software v1.7 on the NovaSeq instrument. Raw sequence data (.bcl files) generated from Illumina NovaSeq was converted into fastq files and de-multiplexed using Illumina bcl2fastq program version 2.20. One mismatch was allowed for index sequence identification.
3. **Transcriptome analyses:** Data processing was performed by GENEWIZ NGS Europe (Azenta Life Sciences). After investigating the quality of the raw data, sequence reads were trimmed to remove possible adapter sequences and nucleotides with poor quality using Trimmomatic v.0.36. The trimmed reads were mapped to the *Homo sapiens* reference genome available on ENSEMBL using the STAR aligner v.2.5.2b. The STAR aligner is a splice aligner that detects splice junctions and incorporates them to help align the entire read sequences. BAM files were generated as a result of this step. Unique gene hit counts were calculated by using feature Counts from the Subread package v.1.5.2. Only unique reads that fell within exon regions were counted. After extraction of gene hit counts, the gene hit counts table was used for downstream differential expression analysis. Using DESeq2, a comparison of gene expression between the groups of samples was performed. The Wald test was used to generate p-values and Log2 fold changes. Volcano plots were generated using the R package ggplot2 v.3.5.1 to visualise differential gene expression between conditions. Gene Ontology (GO) enrichment analysis was performed using the R package clusterProfiler v.4.10.1, with gene annotations from org.Hs.eg.db v.3.18.0. Differentially expressed genes were separated into upregulated and downregulated gene sets prior to enrichment analysis. For selected gene-set analyses, expression values for manually selected genes were extracted across all samples and converted to row-wise z-scores, calculated independently for each gene. The resulting z-score matrix was reshaped into long format and visualised using ggplot2 v.3.5.1 as boxplots with individual gene-level values overlaid as jittered points. Statistical comparison of z-score distributions was used as a descriptive summary of gene-set-level expression changes. Heatmaps were generated using the R package pheatmap v.1.0.12. RNA-seq data for this study will be available through Gene Expression Omnibus (GEO).

### Laser ablations and analysis

Annual laser ablations were performed as described ^24,70^. Briefly, cells were stained with CellMask Deep Red Actin Tracking stain at 37°C for 5 min (1:100, ThermoFisher, A57245). A circular ROI was positioned apically in a region of continuous epithelium. A single apical Z-slice containing the apical surface was selected and imaged for two frames before ablation. Ablation was performed using a Mai Tai laser (Spectra-Physics Mai Tai eHP DeepSee multiphoton laser; 1,436.1 mW maximum power) at 700 nm, with 100% laser power, a pixel dwell time of 0.42 μs and 10 iterations. The instant recoil after ablation was used for quantification. Recoil was measured as the changes in the apical surface area of individual cells. Images were captured on a Zeiss Examiner LSM880 confocal, using a 20x/NA1.0 Plan Apochromat water objective with AiryScan Fast. Images used for analysis were processed using the Fiji/ImageJ-based plugins StackReg and PureDenoise.

### Statistical analysis

Most of the biological experiments were performed with multiple independent biological replicates, which include at least 3 disomy and 3 TS21 iPSC lines, as detailed in the figure legends. For the experiments with less of 3 biological replicates, technical replicates are detailed in manuscript, methods and figure legends. Statistical tests and comparisons between multiple samples are detailed in the figure legends. Statistics were performed and graphs were generated in Prism v10.1.0 (Graphpad) and R version 4.3.2.

