## Supplementary Material for "Extracellular matrix defects destabilise apical cytoarchitecture and mechanical properties during early Down syndrome neurodevelopment"

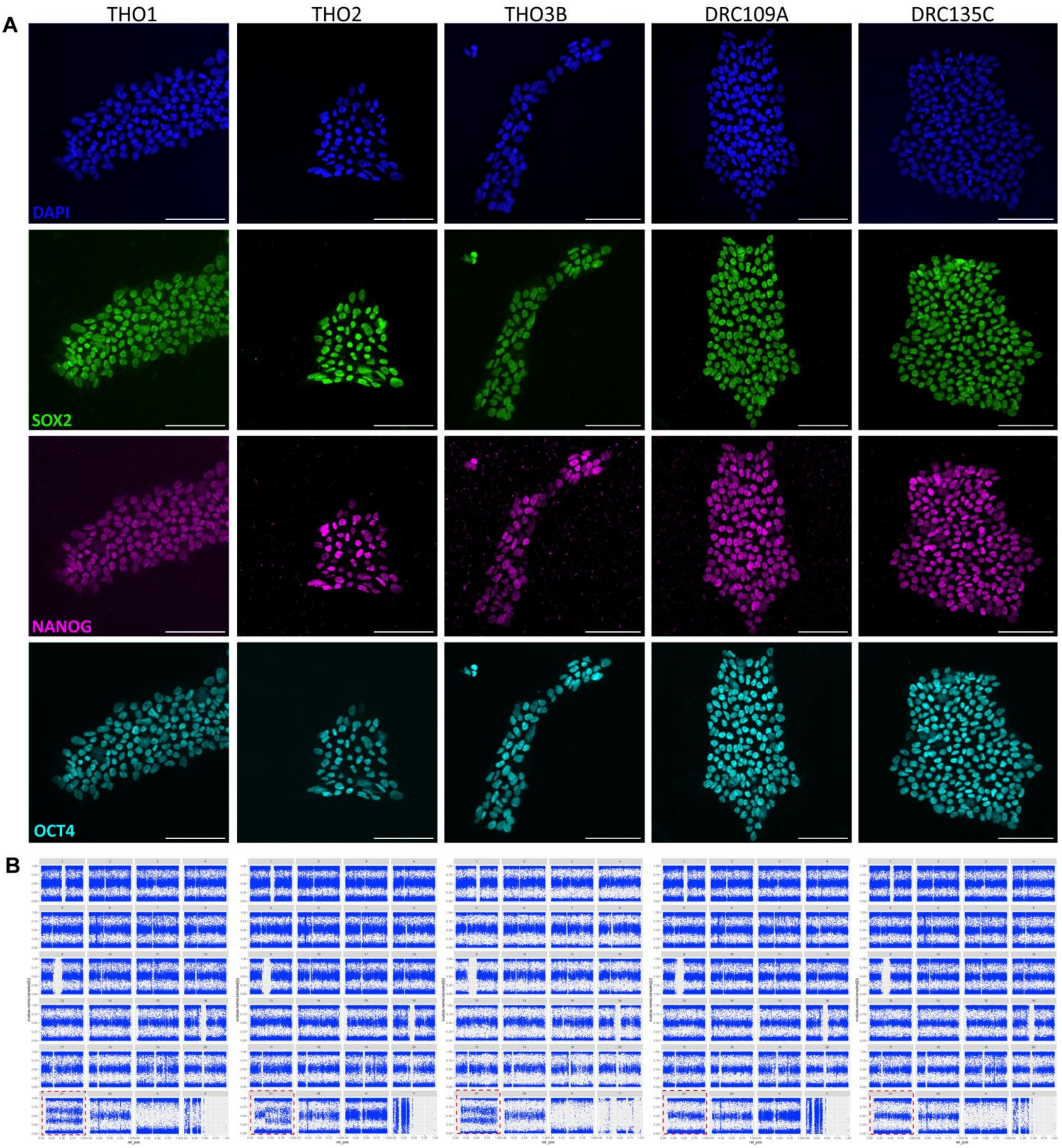


**Supplementary Figure 1. A.** Representative immunofluorescence images of iPSC colonies from THO1, THO2, THO3B, DRC109A and DRC135C stained for the pluripotency-associated transcription factors SOX2, NANOG and OCT4, together with DAPI nuclear counterstaining. Scale bar, 100 µm. **B.** Genome-wide SNP-array/copy-number profiles of the same iPSC lines. Red boxes highlight HSA21.

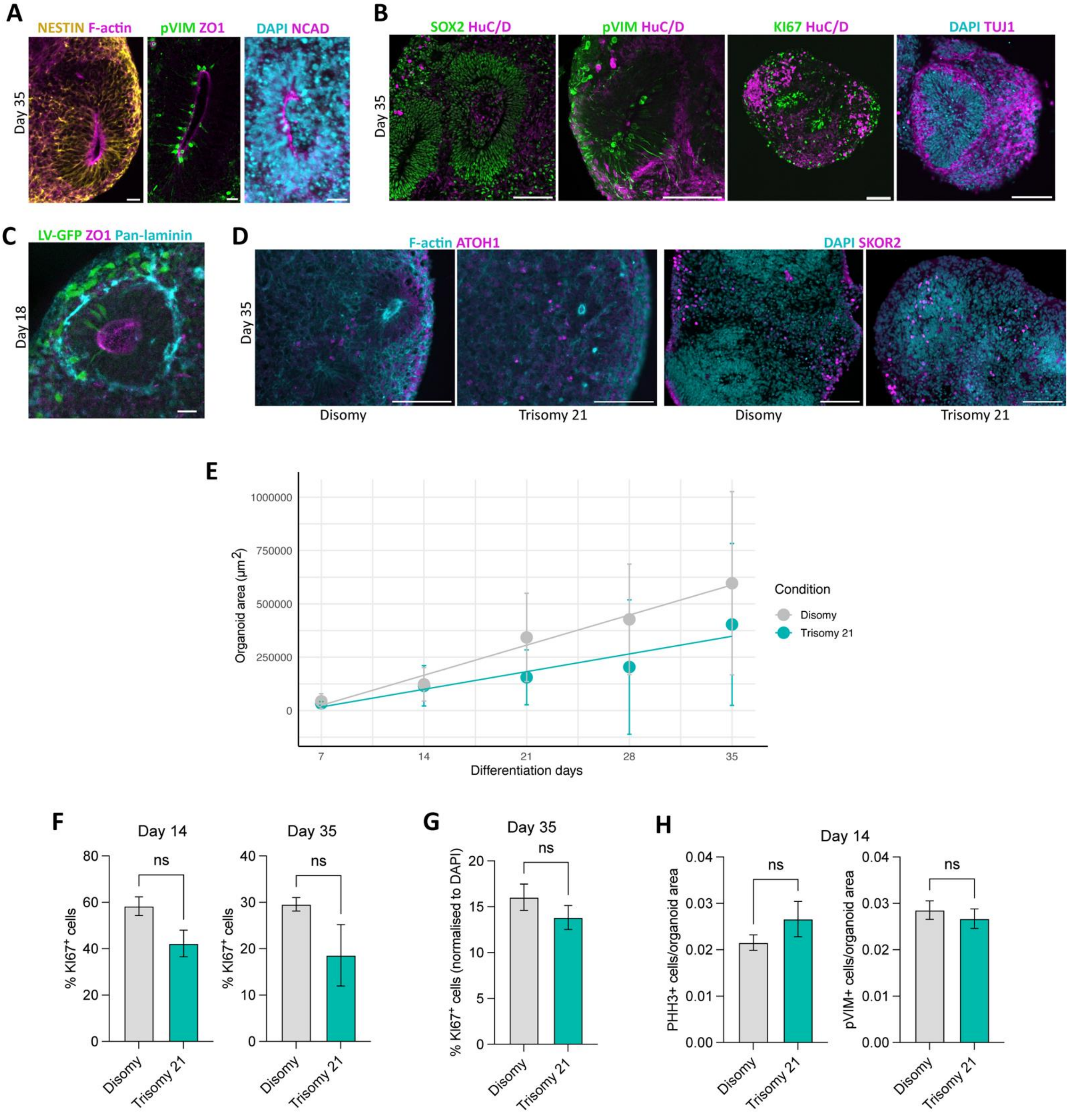


**Supplementary Figure 2. A.** Immunofluorescence staining of NESTIN, and apical F-actin, ZO1 and N-Cadherin (NCAD), exhibiting neural rosettes containing a lumen-like centre. Scale bar, 20 µm. **B.** Immunofluorescence for SOX2, pVIM, KI67, HuC/D, and TUJ1 showing proliferative neural progenitors and neurons on day 35. Scale bar, 100 µm. **C.** Representative immunofluorescence image from day 18 organoid slices showing the morphology of neural progenitor cells labelled with viral GFP, in contact with apical ZO1-expressing lumens, and end-feet touching laminin-expressing basement membrane. Scale bar, 20 µm. **D.** Immunofluorescence for ATOH1 and SKOR2 showing cerebellar progenitors in both disomy and TS21-derived organoids by day 35. Scale bar, 100 µm. **E.** Quantification of organoid area from day 7 to day 35 of differentiation. Organoid area increased significantly over time in both control and TS21 conditions. Control organoids showed a strong positive correlation between area and timepoint ($r = 0.70$, 95% CI = 0.66–0.74, $p < 2.2 \times 10^{-16}$), whereas TS21 organoids showed a weaker, although still significant, positive correlation ($r = 0.45$, 95% CI = 0.38–0.52, $p < 2.2 \times 10^{-16}$). Linear regression analysis showed that time explained a larger proportion of the variation in organoid area in control organoids ($R^2 = 0.49$) than in TS21 organoids ($R^2 = 0.20$). Comparison of correlation coefficients using Fisher's z-transformation revealed that the association between organoid area and time was significantly stronger in control than in TS21 organoids ($z = 6.45$, $p = 1.1 \times 10^{-10}$). Points represent mean ± SEM; lines represent linear regression analysis. **F.** Quantification of $KI67^+$ cells at days 14 and 35, showing no significant differences between disomic and TS21 organoids. Data obtained from 2 disomic and 2 TS21 cell lines. Mann-Whitney test; error bars represent SEM. ns, not significant. **G.** Quantification of $KI67^+$ cells normalised to $DAPI^+$ nuclei at day 35, showing no significant difference between conditions. Data obtained from 2 disomic and 2 TS21 cell lines. Mann-Whitney test; error bars represent SEM. ns, not significant. **H.** Quantification of $PHH3^+$ and $pVIM^+$ cells normalised to organoid area at day 14, indicating no significant differences in mitotic cell populations between disomic and TS21 organoids. Data obtained from 3 disomic and 3 TS21 cell lines. Mann-Whitney test; error bars represent SEM. ns, not significant.

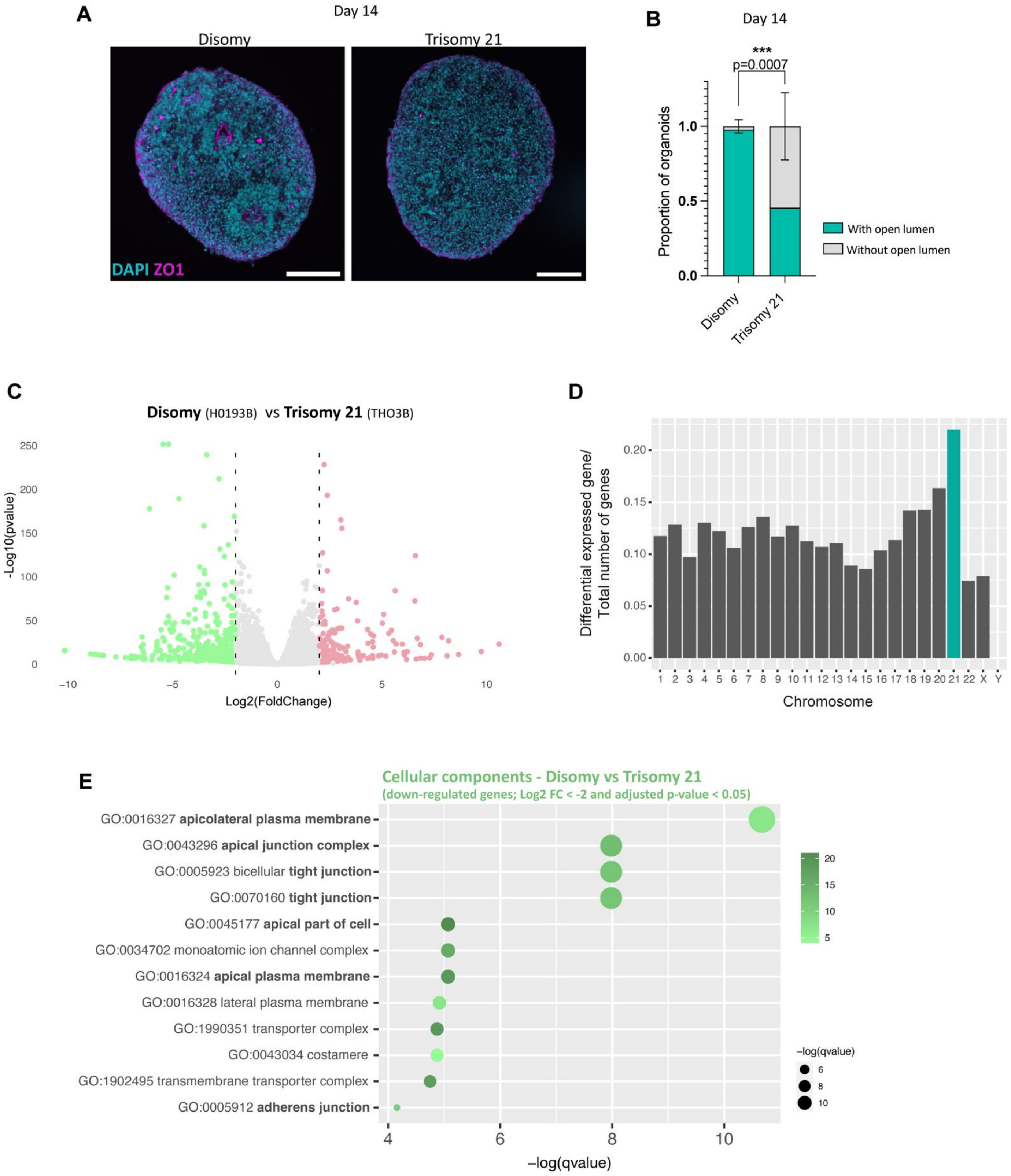


**Supplementary Figure 3. A.** Immunofluorescence staining of organoid cryosections for ZO1 at day 14. Scale bar, 100 µm. **B.** Stacked bar graph shows the proportion of organoids with open lumens. Data obtained from 3 disomic and 3 TS21 cell lines. Two-way ANOVA (Uncorrected Fisher's LSD); error bars represent SEM. ***p = 0.0007. **C.** Volcano plot of genes identified after cerebellar differentiation of human iPSCs on day 14, comparing disomy and TS21 conditions. **D.** Bar-plot graph shows the ratio of differential expressed genes per total number of genes for each chromosome for disomy vs TS21 at day 14 of cerebellar differentiation. HSA21 is highlighted in green. **E.** Top gene ontology (GO) Cellular component terms identified for the differentially down-regulated genes (Log2 FC < -2 and adjusted p-value < 0.05) of DS-derived cerebellar organoids at day 14, demonstrating a significant involvement of apical transcripts.

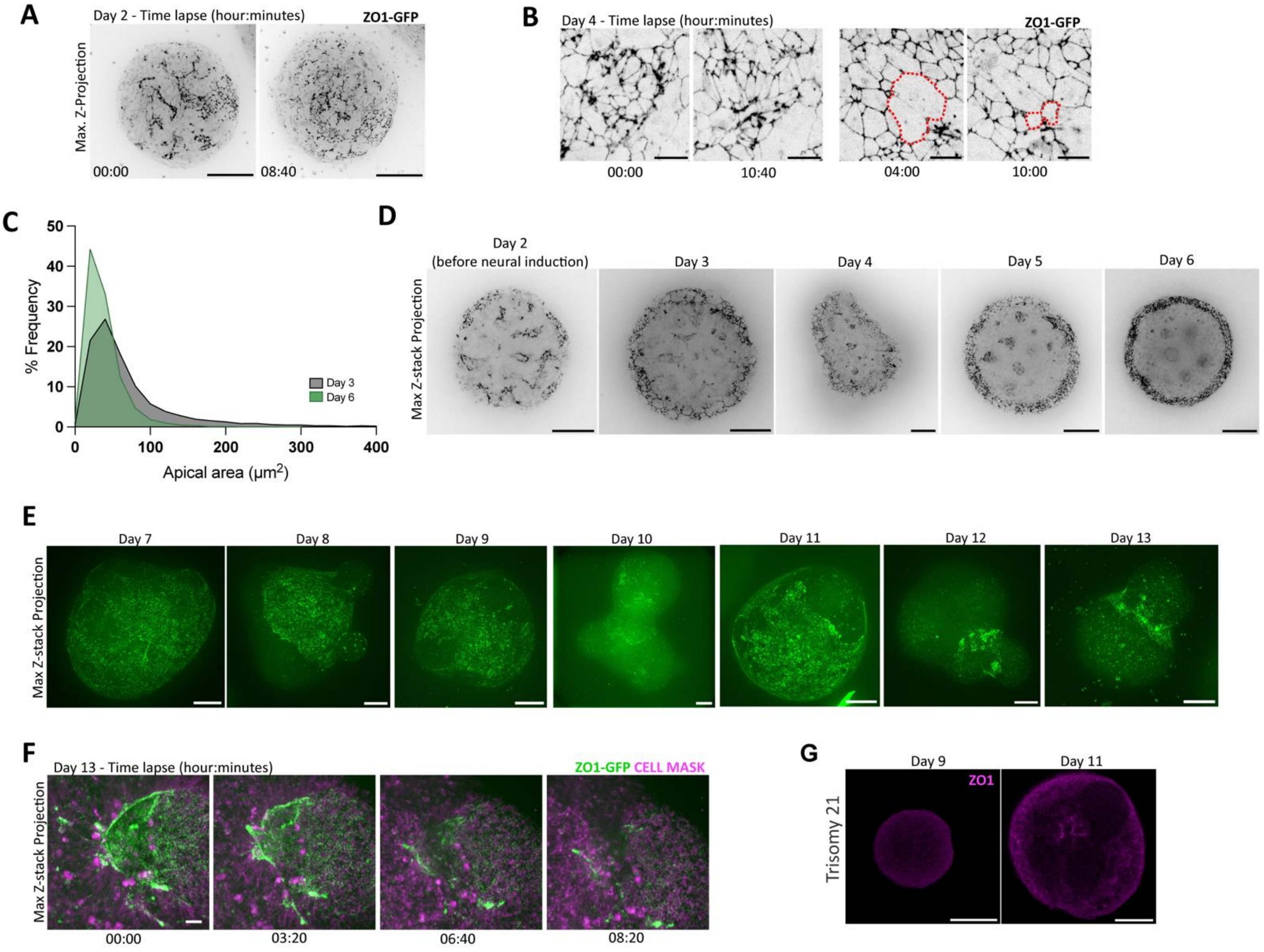


**Supplementary Figure 4. A.** Still frames of live imaging of ZO1-GFP-derived cerebellar organoids at days 2. Scale bar, 100 µm. **B.** Live imaging time-lapses of day 4 ZO1-GFP organoids. Dashed red lines delineate cells undergoing apical constriction. Scale bar, 20 µm. **C.** Frequency distribution showing the percentage of cells according to apical area. Compared with day 3, day 6 shows a narrower distribution shifted towards smaller apical areas, indicating more homogeneous apical surface sizes during neuroepithelial remodelling. Data obtained from ≥ 5 organoids per timepoint. **D.** Representative maximum intensity Z-projection of live imaging images of ZO1-GFP organoids from day 2 to 6 of differentiation, showing lumens inside of organoids. Scale bar, 100 µm. **E.** Live imaging time-lapses of ZO1-GFP organoids from day 7 to 14 of differentiation, showing apical surface breaks. Scale bar, 100 µm. **F.** Live imaging time-lapses of day 13 ZO1-GFP organoids, suggesting apical internalisation. Scale bar, 20µm. **G.** Representative single z-level from whole mount immunofluorescence staining for ZO1, showing organoids without detectable apical lumens in TS21 condition at days 9 and 11. Scale bar, 100µm.

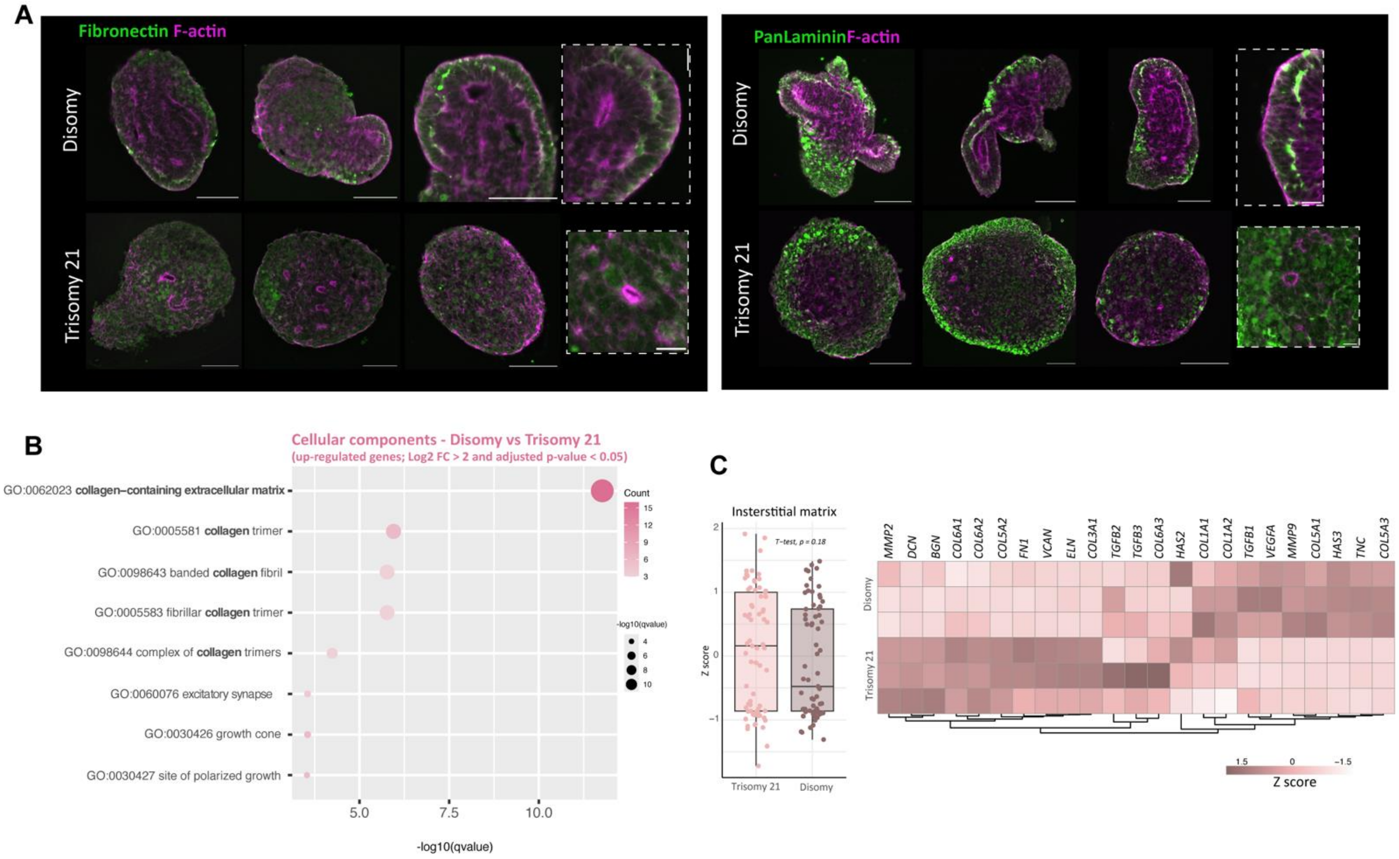


**Supplementary Figure 5. A.** Immunofluorescence images showing the location of different ECM components, including fibronectin and pan-laminin in organoids at day 14. Scale bar, 100 µm. High magnifications of neuroepithelial-like structures are represented in the highlighted by dashed squares. Scale bar, 20 µm. **B.** Top gene ontology (GO) Cellular component terms identified for the differentially down-regulated genes (Log2 FC > 2 and adjusted p-value < 0.05) of DS-derived cerebellar organoids at day 14, demonstrating a significant involvement of collagen-related genes. **C.** Box-blot shows the distribution of interstitial matrix-related genes listed in the heatmap. Data from bulk RNAseq data at day 14. Values are shown as Z-score.

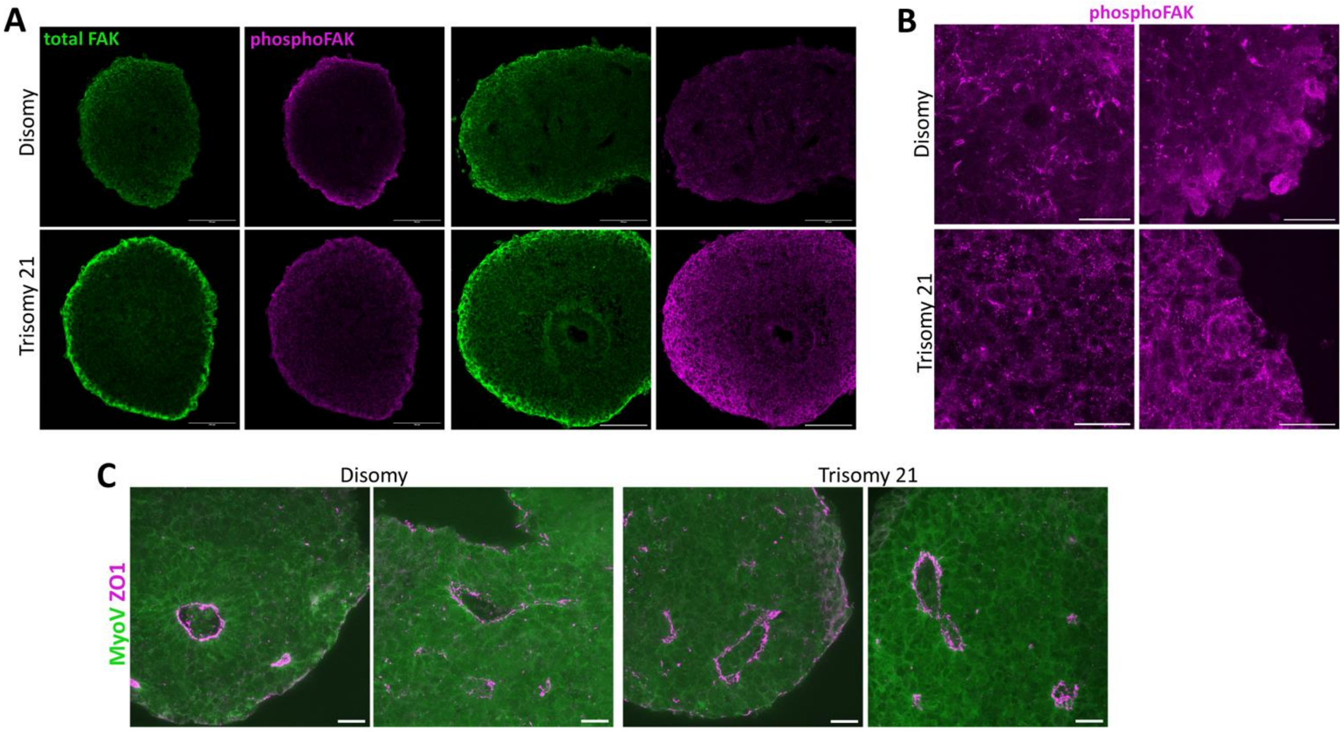


**Supplementary Figure 6. A.** Immunofluorescence staining for total FAK and phosphoFAK in cerebellar organoid cryosections at day 14. Scale bar, 100 µm. **B.** Representative high magnification images of phosphoFAK in cerebellar organoid cryosections at day 14. Scale bar, 20 µm. **C.** Representative staining of organoid cryosections for ZO1 and MyoV at day 14. Scale bar, 20 µm.

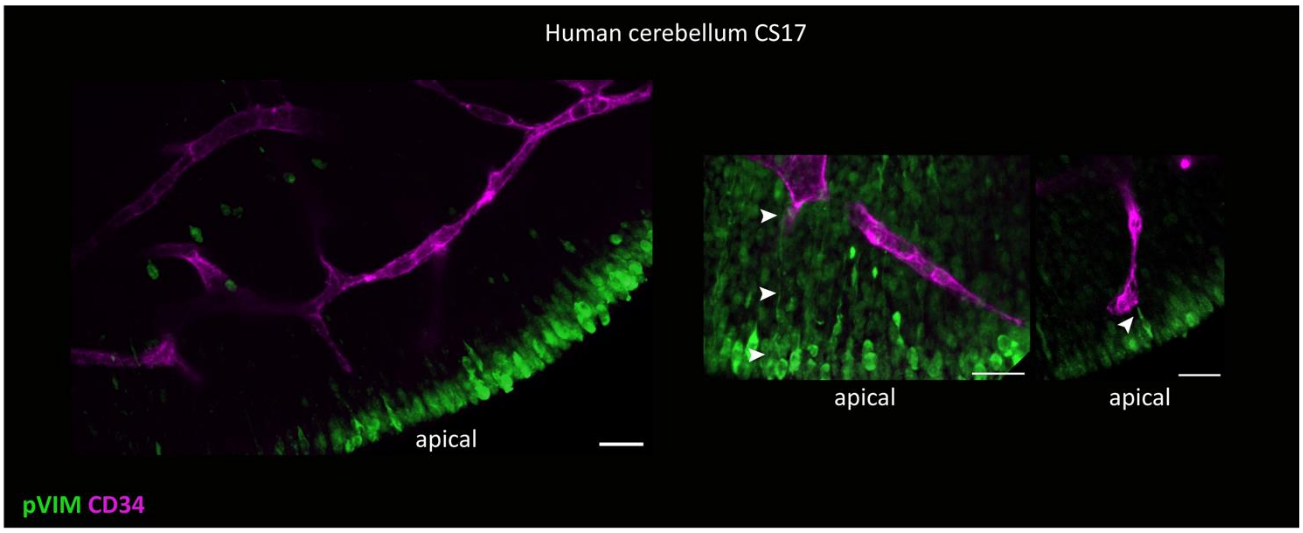


**Supplementary Figure 7.** Immunofluorescence for pVIM and CD34 showing proliferative neural progenitors and blood vessels, respectively. The arrowheads indicate pVIM expressing progenitors interacting with blood vessels. Scale bar, 20 µm.

**Supplementary Table S1.** Detailed information of human iPSC lines.

| Cell line name | Source | Karyotype | Reprogramming method |
|---|---|---|---|
| HO193B, "GOC1" [1–3] | NIHR GOSH BRC* | 46, XY | Microfluidic mRNAs |
| DRC135C [4,5] | Livesey Lab | 46, XY | Non-integrating Sendai virus |
| DRC109A[4,5] | Livesey Lab | 46, XX | Non-integrating Sendai virus |
| SFC086-03-01 [1,2] (STBCi052-A) | StemBANCC consortium (EBiSC) | 46, XX | Non-integrating Sendai virus |
| ZO-1 mEGFP (mEGFP-TJP1-cl20) | Allen Cell Collection | 46, XY | Integration-free episomal vectors |
| THO1 (TS21-1C [5]) | Livesey Lab | 47, XY, +21 | Non-integrating Sendai virus |
| THO2 (TS21-2A [5]) | Livesey Lab | 46,XX,dup(21)(q21.2q22.3) | Non-integrating Sendai virus |
| THO3B (TS21-3B [5]; Ts21-2 [4]) | Livesey Lab | 47, XY, +21 | Non-integrating Sendai virus |

* National Institute of Health Research Great Ormond Street Hospital Biomedical Research Centre

**Supplementary Table S2.** List of primary antibodies.

| Antibody | Source | Dilution | Catalogue number |
|---|---|---|---|
| Mouse anti-ZO1 | ThermoFisher | 1:100 | ZO1-1A12 |
| Mouse anti-NESTIN | R&D Systems | 1:200 | MAB1259 |
| Rabbit anti-pVIM | Abcam | 1:100 | ab217673 |
| Mouse anti-NCAD | Cell Signaling | 1:100 | 14215 |
| Rat anti-SOX2 | ThermoFisher | 1:50 | 740013T |
| Mouse anti-HuC/D | ThermoFisher | 1:400 | A21271 |
| Rabbit anti-KI67 | ThermoFisher | 1:100 | MA5-14520 |
| Mouse anti-TUJ1 | BioLegend | 1:500 | 801213 |
| Rabbit anti-Pan-Laminin | Abcam | 1:200 | ab11575 |
| Rabbit anti-Fibronectin | Abcam | 1:200 | ab23750 |
| Goat anti-Collagen IV | Merk | 1:50 | AB769 |
| Rabbit anti-PAR3 | Merk Millipore | 1:100 | 07-330 |
| Mouse anti-β-catenin (8E7 active) | Upstate | 1:100 | 05-660 |
| Rabbit anti-FAK | Santacruz | 1:100 | sc-558 |
| Mouse anti-phosphoFAK | Merck | 1:100 | 05-1140 |
| Rabbit anti-Myo5A | Cell Signaling | 1:100 | 3402S |
| Rabbit Anti-RAB11 | ThermoFisher | 1:100 | 3H18L5 |

**Supplementary Table S3.** List of secondary antibodies.

| Antibody | Source | Dilution | Catalogue number |
|---|---|---|---|
| Goat-anti rabbit 488 | | | A11034 |
| Goat anti-mouse 568 | | | A11031 |
| Goat anti-mouse IgG1 (y1) 568 | | | A21124 |
| Goat anti-mouse IgG2b (y2b) 633 | | | A21146 |
| Donkey anti-goat 488 | ThermoFisher | 1:500 | A11055 |
| Donkey anti-mouse 568 | | | A10037 |
| Donkey anti-rabbit 647 | | | A31573 |
| Donkey anti-rat 488 | | | A21208 |
| Donkey anti-rat 647 | | | A78947 |

**<u>Methods</u>:**

**Flow Cytometry:** Organoids were dissociated to single cells with accutase for 7 min at 37$^\circ$C. Enzyme was inactivated by addition of serum-containing medium and centrifuged at 200 x g for 5 min. After centrifugation and washed once with PBS, the cell pellet was fixed drop by drop with 70% v/v ethanol (previously stored at −20$^\circ$C) and stored at −20$^\circ$C. For staining, samples were centrifuged at 200 x g for 10 min. Then, cells were washed twice with PBS. For each experiment, approximately 500,000 cells were resuspended in KI67 antibody (DAKO, M7240, 1:100) diluted in 3% v/v BSA solution in PBS and incubated for 30 min at RT. Then, cells were washed with PBS, resuspended in 3% v/v BSA solution in PBS and incubated with secondary antibodies, goat anti-mouse 488 (ThermoFisher, A28175, 1:500) for 15 min at RT. Finally, cells were washed twice with PBS, resuspended in PBS and analysed in a BD® LSR II Flow Cytometer (BD Biosciences).
